# Zooplankton community beta diversity in an Amazonian floodplain lake

**DOI:** 10.1101/2020.07.31.231241

**Authors:** Leonardo Fernandes Gomes, Ana Caroline Alcântara Missias Gomes, Carla Albuquerque de Souza, Hasley Rodrigo Pereira, Marie-Paule Bonnet, Ludgero Cardoso Galli Vieira

**Affiliations:** Núcleo de Estudos e Pesquisas Ambientais e Limnológicas - NEPAL, Faculdade UnB de Planaltina, Área Universitária 1, Vila Nossa Senhora de Fátima, 73.345-010 – Planaltina – DF, Brasil; Geosciences Environnement Toulouse (UMR 5563 GET), IRD/ CNRS/ Université Toulouse III, Toulouse, France; International Joint Laboratory Observatoire des Changements Environnementaux LMI OCE, IRD – Universidade de Brasília, Campus Universitário Darcy Ribeiro, CEP 70910-900 Brasília, Brazil

**Keywords:** Lago Grande do Curuai, hydrological cycle, Podani, flood pulse, beta diversity partitioning

## Abstract

Understanding the mechanisms that generate organism distribution patterns from the beta diversity perspective can assist in environmental monitoring strategies. In this study, we emphasized the limnic zooplankton due to the ability of these organisms to respond quickly to environmental variations. Therefore, we evaluated the following questions: (i) Do different regions of the same lake have the same importance in contributing to beta diversity? (ii) Do beta diversity and its components vary over the hydrological cycle? (iii) What is the importance of local and spatial predictors in beta diversity and its components? (iv) Do beta diversity and its components show a consistent pattern throughout the hydrological cycle? We found that the contribution of different sites to diversity was more associated with regions with low abundance and richness of organisms values, such as the littoral and *igarapes*, which shows the relevance of these areas for biological monitoring and for the delimitation of priority areas for the zooplankton diversity conservation. Despite the peculiarities of each hydrological period and regarding beta diversity components, we verified a species substitution and differences in abundance pattern in the lake. We also found low concordance patterns between the periods and low environmental and spatial variables prediction on beta diversity patterns.

## Introduction

Species can present different distribution patterns in response to natural factors such as competition, predation, dispersive processes limitations, and/or local and regional environmental variables influences (Guisan & Thuiller, 2005). These factors may be intensified by human activities, which makes even more relevant to understand the mechanisms that generate such structuring patterns in biological communities. Thus, the understanding of these mechanisms can assist in the formulation of efficient environmental monitoring strategies and, even, in the delimitation of priority areas for conservation in several ecosystems (Socolar *et al*., 2016). The comparative diversity across multiple sites, known as beta diversity (Whittaker, 1960), has undergone advances over the years both for understanding patterns of presence-absence of organisms and for density values per site (Baselga, 2010; Podani & Schmera, 2011; Podani, Ricotta & Schmera, 2013).

Both for organism occurrence and abundance, Podani family of beta diversity (Podani & Schmera, 2011; Podani *et al*., 2013) can be partitioned into the following main components: (i) *species similarities*: commonly measured by the Jaccard index for presence-absence data and Ruzicka, for abundance data. High values of this partition mean that the pairs of sites put in comparison share many species or species with similar abundances; (ii) *difference in relative richness/abundance*: is the difference in species richness, or species abundance, between pairs of sites. Therefore, high values of that partition show that the number of species or specimens between compared sites is discrepant; (iii) *species replacement/abundance:* it can be maximized when there is a high replacement of species, or species with equivalent abundances, along an environmental gradient or between pairs of sites. Therefore, high replacement values for the abundance data mean that, although the sites in comparison have similar abundances, the species composition is different. Also, although the approach with abundance data may represent more subtle differences concerning environmental variations, the values between the assessments for abundance and presence-absence data can be quite different, even if evaluated with the same data set (Podani *et al*., 2013).

The evaluation of the factors that influence beta diversity and its components can be even more complex in floodplain lakes since they are predominantly dominated by the flood pulse that controls the dynamics of entry and output of sediments, water and organisms that naturally contribute for changes in biological diversity in the ecosystem (Junk *et al*., 2012). These plains are continuously or periodically flooded by direct precipitation or by the overflow of the main river and, depending on the level of connectivity with the river, there may be a loss of connection between habitats during periods of low water (Thomaz, Bini & Bozelli, 2007). However, as the cycle of extensive floodplains is usually slow and monomodal, the biological dynamics of organisms can adapt in order to maximize their performance according to hydrological cycles (Junk *et al*., 2011).

In Amazonian rivers, the flow tends to be more intense and requires a high resilience capacity of the organisms. Therefore, smaller aquatic organisms tend to be present with greater richness and density in the lakes of these plains, where they can find shelter against predation and food (Junk, Bayley & Sparks, 1989). Furthermore, according to the hydrological period, these organisms may present beta diversity patterns that change over time (Bozelli *et al*., 2015).

Assessing beta diversity and its components over space, but also highlighting whether the pattern generated is consistent throughout the hydrological cycle is important in different aspects. For example, due to the scarcity of financial resources and time allocated in environmental monitoring programs and scientific research, if different hydrological periods show a concordant pattern of diversity, there is a real possibility of adjustment in the sampling effort, reducing the number of sampling campaigns, which would save financial resources and time. In the same way, it is possible to use other alternatives as is the case of using lower taxonomic resolutions and or presence-absence data instead of abundance data (Carneiro *et al*., 2013; Vieira *et al*., 2017; de Morais *et al*., 2018).

In this study, we emphasized the limnic zooplankton due to the ability of these organisms to respond quickly to environmental variations. Therefore, we evaluated the following questions: (i) Do different regions of the same lake have the same importance in contributing to beta diversity? (ii) Do beta diversity and its components vary over the hydrological cycle? (iii) What is the importance of local (environmental characterization) and spatial (dispersive processes) predictors in beta diversity and its components? (iv) Do beta diversity and its components show a consistent pattern throughout the hydrological cycle? Taking into account that the ecological dynamics of floodplains is temporally complex, we expected that the sites contribution to beta diversity would be different between hydrological periods. Besides, due to the spatial extent of the study area, we expected that species replacement patterns would be predominant, considering values of presence-absence of organisms, and patterns of differences in abundance, considering values of species abundance per site. Also, due to the complex interactions that dominate the occurrence of organisms, we expected that there would be a variation between environmental and spatial predictors in biological diversity patterns and, finally, as each period comprises a different hydrological dynamics, we did not expect to find many concordant values, being important to evalute in all hydrological periods to understand the distribution patterns of the zooplankton community.

## Material and methods

### Study area

The study area comprises an Amazonian floodplain lake called *Lago Grande do Curuai*, located in the State of Pará, Brazil. The majority of the water supply comes from the Amazon River (77%), while the others are subdivided between rainfall, runoff, and outcropping of groundwater (Bonnet *et al*., 2008). The hydrological dynamics generate a monomodal cycle in this lake, comprising the periods of flooding (from January to the end of February), high water (from April to the end of June), flushing (from August to October) and low water (mid-October to November) (de Moraes Novo *et al*., 2006).

The environmental characteristics of Lago Grande do Curuai are quite variable throughout the year, mainly concerning chlorophyll-*a* and dissolved oxygen. During the flooding period, chlorophyll-*a* levels are low enough for human consumption. However, the values in the flushing period rise to such an extent that water is not recommended for any type of activity (Affonso, Barbosa & Novo, 2011).

Sampling were carried out in 17 sample units (Fig. 1) in four campaigns: March / 2013 (flooding period), September / 2013 (flushing period), May / 2014 (high water period) and November / 2014 (low water period).

**Fig. 1.**
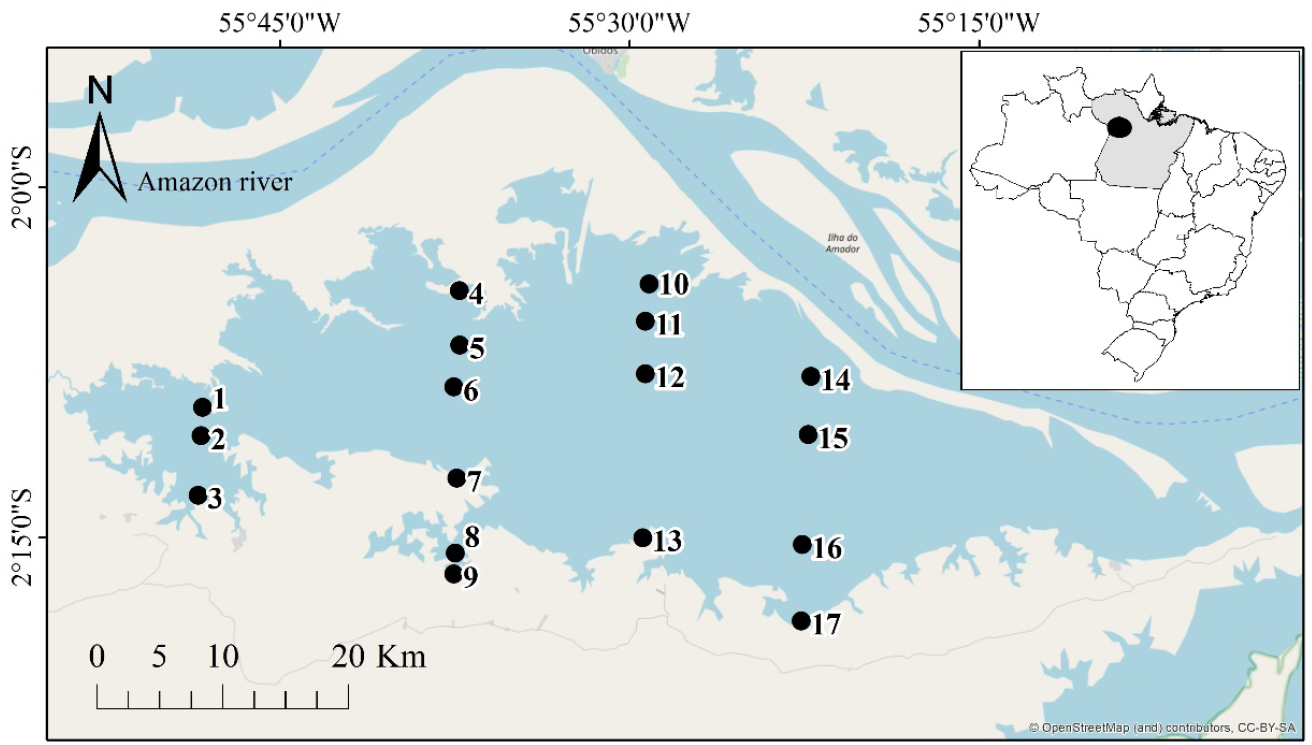
Map of the study area and sampling units in Lago Grande do Curuai. Blue area: aquatic environments; beige area: terrestrial environments

### Environmental variables

In each sampling unit, we used a multi-parameter YSY probe, model EXO2 to measure the variables dissolved oxygen (mg/L), blue-green algae (μg/L), fluorescent organic dissolved matter (raw), pH, water temperature (°C), conductivity (μS/cm), total dissolved solids (mg/L), and turbidity (NTU). According to the protocol (APHA, 2005), water samples were obtained and frozen for further quantification in the laboratory of: alkalinity (mg/L), total chlorophyll (μg/L), total phosphorus (μg/L), total nitrogen (μg/L), total dissolved nitrogen (mg/L), ammonia (mg/L), nitrate (mg/L), and silica (mg/L).

### Zooplankton

In each sampling unit, we sampled the zooplankton community on the subsurface (*ca* 50 cm). Therefore, we filtered 300 liters of water in a net with a 68 μm opening mesh. Samples were stored in polyethylene bottles, preserved with formaldehyde (5%), and buffered with sodium tetraborate. In the laboratory, the samples were concentrated in 75 mL. To quantify the densities of zooplanktonic organisms per sample unit, a 7.5 mL subsampling was performed with a *Hensen-Stempel* pipette. We read the subsampled organisms in a Sedgewick Rafter chamber for identification and counting using an optical microscope. Additionally, we carried out qualitative sampling to verify and record the existence of new *taxa* that were not identified during quantitative sampling (Bottrell *et al*., 1976).

### Data analysis

We performed a Local Contribution to Beta Diversity (LCBD) (Borcard, Gillet & Legendre, 2018) to obtain the degree of exclusivity of the sites in the species composition in each hydrological period using the function *beta*.*div*, package *adespatial* (Dray *et al*., 2018). To evaluate and partition Podani family beta diversity by sample period, we used the function *beta*.*div*.*comp* of *adespatial* package (Dray *et al*., 2018). In both cases, we used the *Jaccard* index for presence and absence values and *Ruzicka* for organism density data.

To verify if there were significant differences in the values resulting from the beta diversity partitioning by period, we performed a Permutational Multivariate Analysis of Variance Using Distance Matrices (PERMANOVA). We obtained these matrices using the *beta*.*div*.*comp* function for both create a matrix encompassing all periods and generate matrices by pairs of periods. For PERMANOVA, we use the *adonis2* function of the *vegan* package (Oksanen *et al*., 2016) and the matrices resulted from the partition were inserted in response to hydrological periods. Additionally, we constructed triangular plots (simplex) to check the distributions of the pairs of sites concerning the partitive components of beta diversity for both *Ruzicka* distance matrices and *Jaccard* in each hydrological periods.

To assess the influence of environmental and spatial variables in the beta diversity partitions of zooplankton community by hydrological period, we performed Distance-Based Redundancy Analysis (dbRDA’s) (Legendre & Andersson, 1999) with different matrices resulted from the beta diversity partitioning (as response variables) and different environmental and spatial variables (as predictor variables). To determine which variables would be inserted in the dbRDA, we performed the analysis of variation inflation factor (VIF) (Borcard *et al*., 2018), removing the environmental variables that showed high collinearity in each sample period (VIF values greater than 20). To determine the spatial predictors (geographic coordinates), we first converted the coordinates to Cartesian distances using the *geoXY* function of the *SoDA* package (Chambers, 2013). Then, we ordered the variables in a Distance-Based Moran’s Eigenvector Maps (dbMEM) (Dray, Legendre & Peres-Neto, 2006; Legendre & Legendre, 2012) using the dbmem function of the adespatial package (Dray *et al*., 2018).

To evaluate the temporal concordance in the distribution patterns of the different zooplankton community beta diversity partitions between hydrological periods, we performed *Procrustes* tests (Gower, 1975). For that, we ordered the matrices resulting from the beta diversity partitioning in different Non-metric multidimensional scaling (NMDS), then we extracted the values from the ordering scores and inserted them into the *protest* function, from *vegan* package (Oksanen *et al*., 2013). To check the significance, 9999 permutations were performed.

For all the mentioned analyzes, we used the statistical software R (R Core Team, 2016).

## Results

Regarding the contribution of sites to beta diversity (LCBD) using presence-absence data of the zooplankton community, only the hydrological periods of flooding and low water presented sites with significant contributions, with site 9 being important for the beta diversity in both periods (Fig. 2). All significant sites (8, 9, and 13) are located in the southern region of the lake. When we evaluated the LCBD using abundance data (Fig. 3), the four periods presented significant sample units. In the flooding and flushing periods, the significant sampling units were located in the north region of the lake (sites 14 and 10, respectively); in the high waters, they were located in the south, and in the low water period they were located in the west region of the lake.

**Fig. 2.**
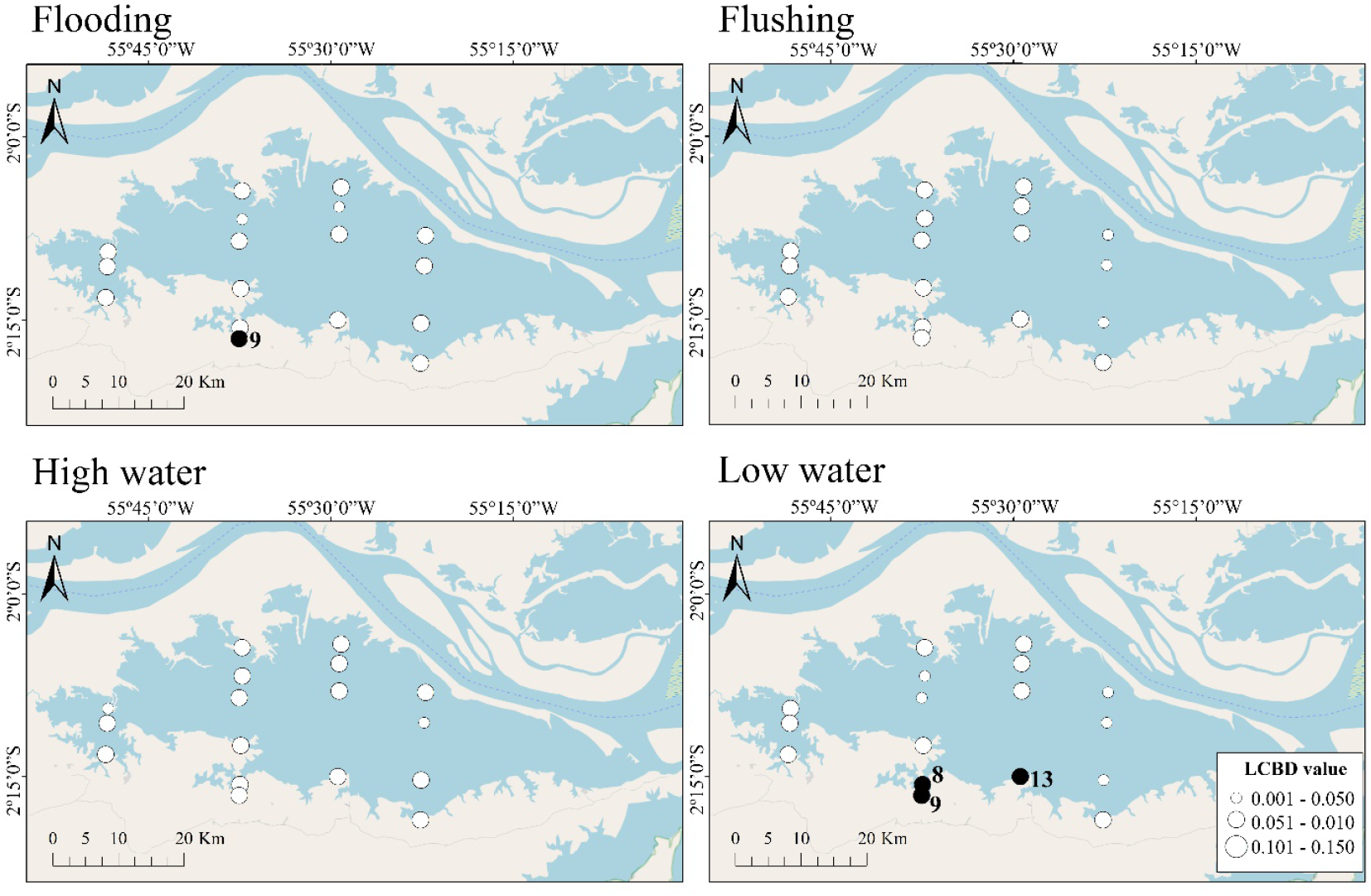
Map of the local contribution to beta diversity (LCBD) for zooplankton presence/absence data with *Jaccard* matrix of the sample units by hydrological period. Filled circles represent sites with significant contributions

**Fig. 3.**
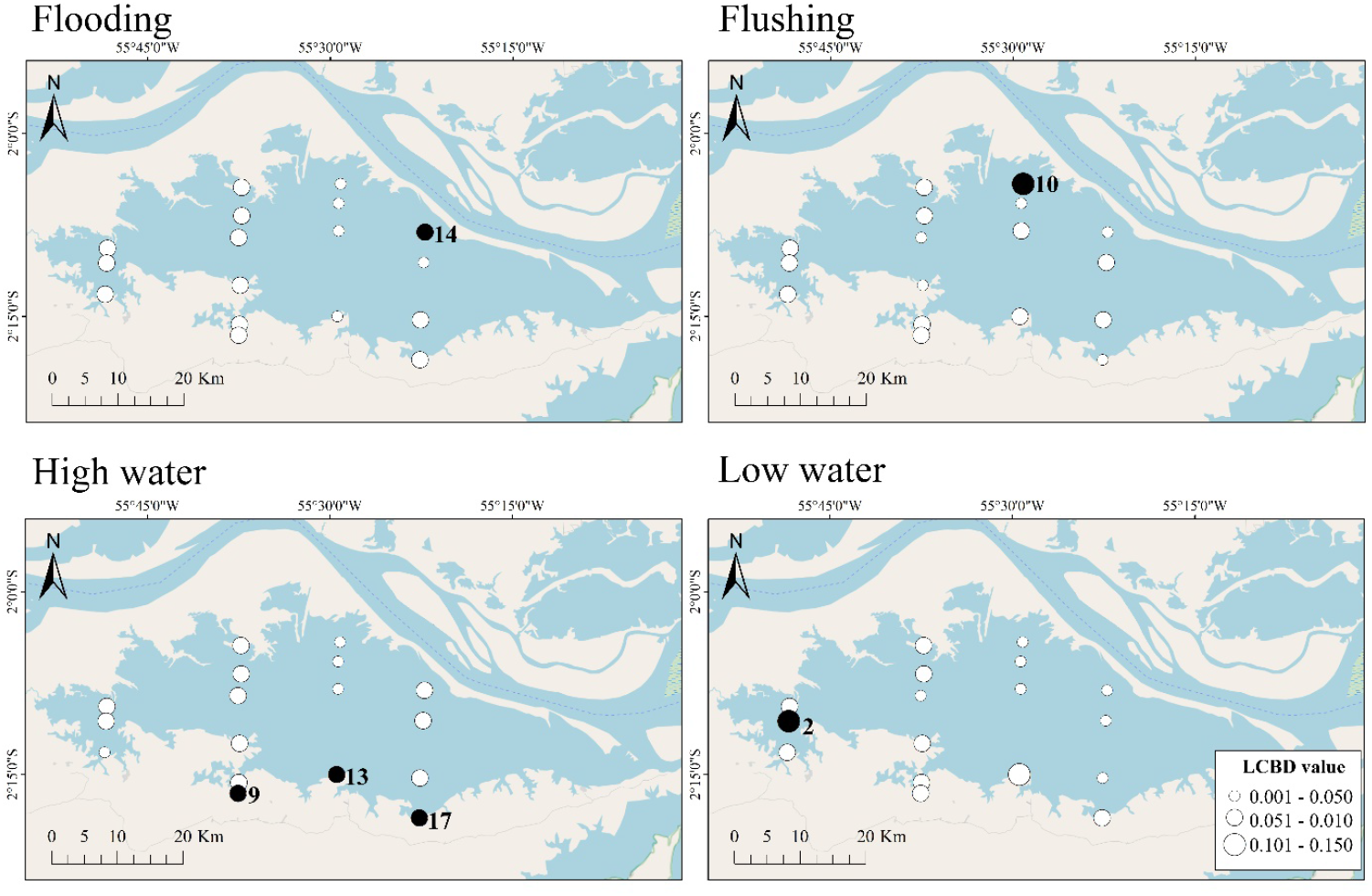
Map of the local contribution to beta diversity (LCBD) for zooplankton abundance data with Ruzicka matrix of the sample units by hydrological period. Filled circles represent sites with significant contributions

Evaluating the beta diversity partitions using presence and absence species data (Table 1), we verified a replacement dominant pattern (values comprised 73% to 81% of the beta diversity between hydrological periods), while we verified an abundance difference dominance pattern when using abundance data (values comprised 58% to 74% of the beta diversity between hydrological periods).

**Table 1.**
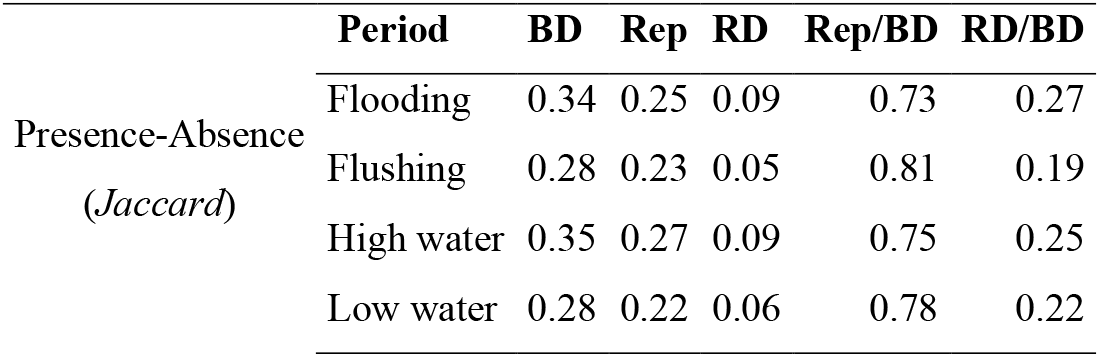
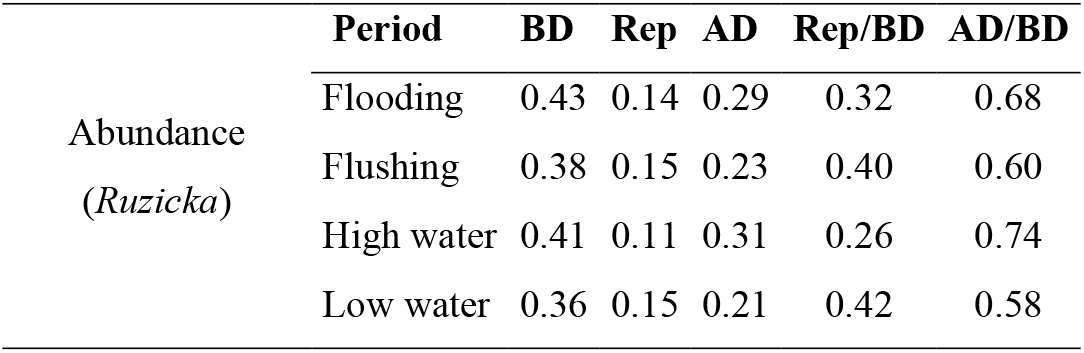
Beta diversity partitioning for all hydrological periods with presence and absence and abundance values. BD = total beta diversity; Rep = replacement; RD = richness difference; AD = abundance difference; Rep/BD = ratio of replacement to total beta diversity; RD/BD = ratio of richness difference to total beta diversity; AD/BD = ratio of abundance difference to total beta diversity

|  | Period | BD | Rep | RD | Rep/BD | RD/BD |
| --- | --- | --- | --- | --- | --- | --- |
| Presence-Absence<br>(Jaccard) | Flooding | 0.34 | 0.25 | 0.09 | 0.73 | 0.27 |
|  | Flushing | 0.28 | 0.23 | 0.05 | 0.81 | 0.19 |
|  | High water | 0.35 | 0.27 | 0.09 | 0.75 | 0.25 |
|  | Low water | 0.28 | 0.22 | 0.06 | 0.78 | 0.22 |

|  | Period | BD | Rep | AD | Rep/BD | AD/BD |
| --- | --- | --- | --- | --- | --- | --- |
| Abundance<br>(*Ruzicka*) | Flooding | 0.43 | 0.14 | 0.29 | 0.32 | 0.68 |
|  | Flushing | 0.38 | 0.15 | 0.23 | 0.40 | 0.60 |
|  | High water | 0.41 | 0.11 | 0.31 | 0.26 | 0.74 |
|  | Low water | 0.36 | 0.15 | 0.21 | 0.42 | 0.58 |

When we compared the beta diversity partitions obtained by the hydrological periods (Table 2) using presence/absence data, the richness difference component was similar among all hydrological periods, while the beta diversity and replacement component were different among them all. When considering abundance data, the beta diversity component was different across all hydrological periods, while the abundance difference component only was not different in flushing and low water periods. There were no differences in the abundance replacement component.

**Table 2.** Permutational multivariate analysis of variance using distance matrices (PERMANOVA) between the matrices resulting from the partition of the beta diversity for the different hydrological periods. Significant values are in bold

|  | Period | Beta diversity (Jaccard) |  |  | Richness difference |  |  | Replacement |  |  |
| --- | --- | --- | --- | --- | --- | --- | --- | --- | --- | --- |
| | | $R^2$ | $F$ | $p$ | $R^2$ | $F$ | $p$ | $R^2$ | $F$ | $p$ |
| Presence-absence | Global | **0.31** | **9.78** | **0.001** | -0.04 | -0.79 | 1.000 | **0.38** | **12.84** | **0.001** |
|  | Flooding x Flushing | **0.21** | **8.58** | **0.001** | - | - | - | **0.27** | **11.90** | **0.001** |
|  | Flooding x High water | **0.18** | **7.15** | **0.001** | - | - | - | **0.22** | **9.27** | **0.001** |
|  | Flooding x Low water | **0.25** | **10.66** | **0.001** | - | - | - | **0.32** | **15.26** | **0.001** |
|  | Flushing x High water | **0.24** | **9.87** | **0.001** | - | - | - | **0.27** | **12.01** | **0.001** |
|  | Flushing x Low water | **0.25** | **10.43** | **0.001** | - | - | - | **0.30** | **13.66** | **0.001** |
|  | High water x Low water | **0.28** | **12.49** | **0.001** | - | - | - | **0.33** | **15.50** | **0.001** |
|  | Period | Beta diversity (Ruzicka) |  |  | Abundance difference |  |  | Replacement |  |  |
| | | $R^2$ | $F$ | $p$ | $R^2$ | $F$ | $p$ | $R^2$ | $F$ | $p$ |
| Abundance | Global | **0.20** | **5.32** | **0.001** | **0.27** | **7.92** | **0.001** | 0.10 | 2.48 | 0.077 |
|  | Flooding x Flushing | **0.12** | **4.52** | **0.001** | **0.12** | **4.27** | **0.012** | - | - | - |
|  | Flooding x High water | **0.10** | **3.63** | **0.001** | **0.14** | **5.35** | **0.004** | - | - | - |
|  | Flooding x Low water | **0.13** | **4.66** | **0.001** | **0.09** | **3.19** | **0.035** | - | - | - |
|  | Flushing x High water | **0.17** | **6.70** | **0.001** | **0.33** | **16.04** | **0.001** | - | - | - |
|  | Flushing x Low water | **0.13** | **4.81** | **0.001** | -0.01 | -0.27 | 0.997 | - | - | - |
|  | High water x Low water | **0.20** | **7.88** | **0.001** | **0.34** | **16.66** | **0.001** | - | - | - |

In proportion, when we partitioned the beta diversity using presence-absence data, the pairs of sample units were more associated with greater similarities and replacement values considering all periods (Fig. 4). On the other hand, when we evaluated the partition using abundance data, the pairs of sample units were more associated with abundance difference component and, secondly, with higher replacement levels (Fig. 5).

**Fig. 4.**
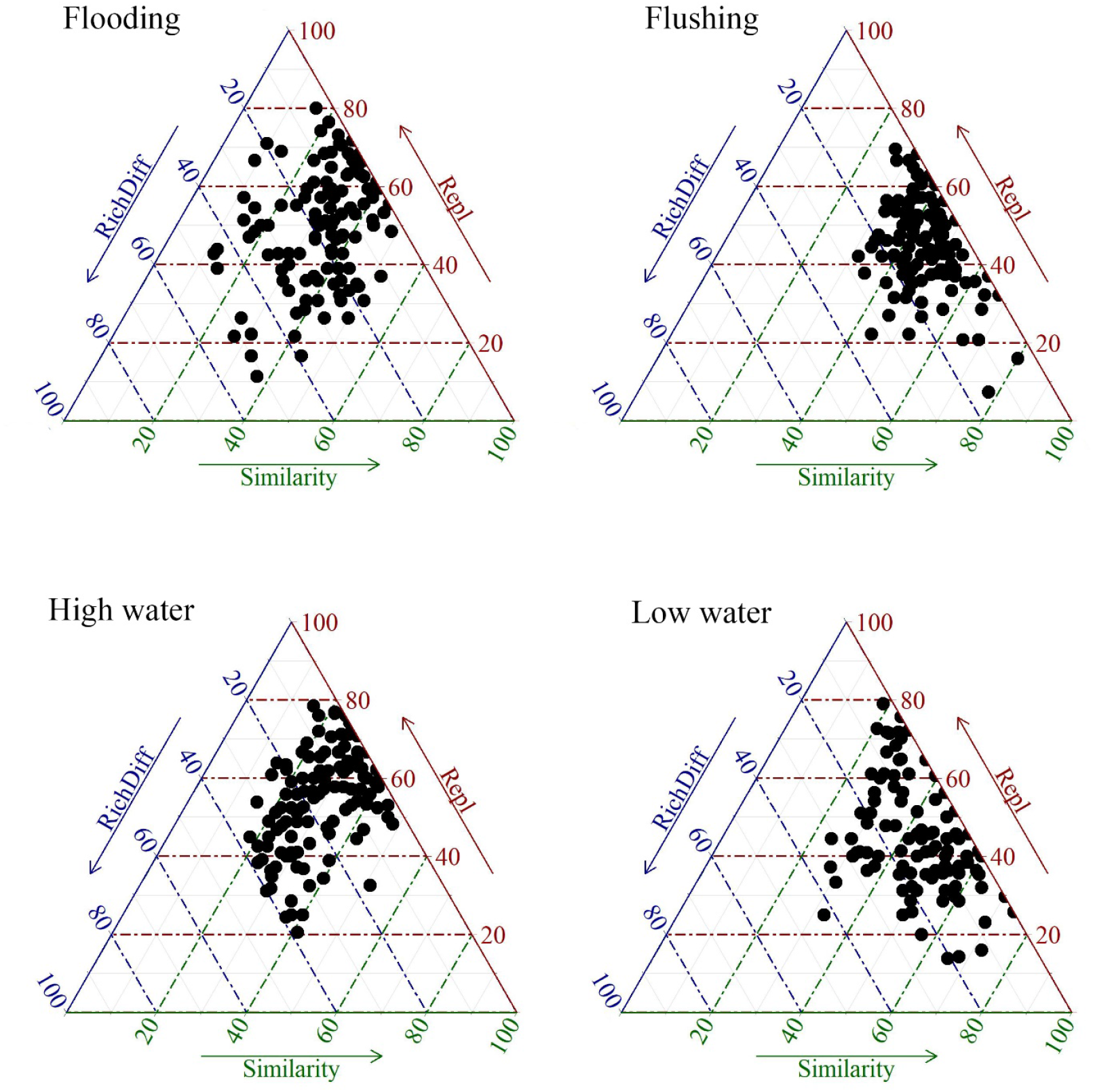
Triangular graph (simplex) of the proportion of elements of the beta diversity partition per pair of sample units for values of presence-absence of organisms. RichDiff = richness difference and Repl= species replacement.

**Fig. 5.**
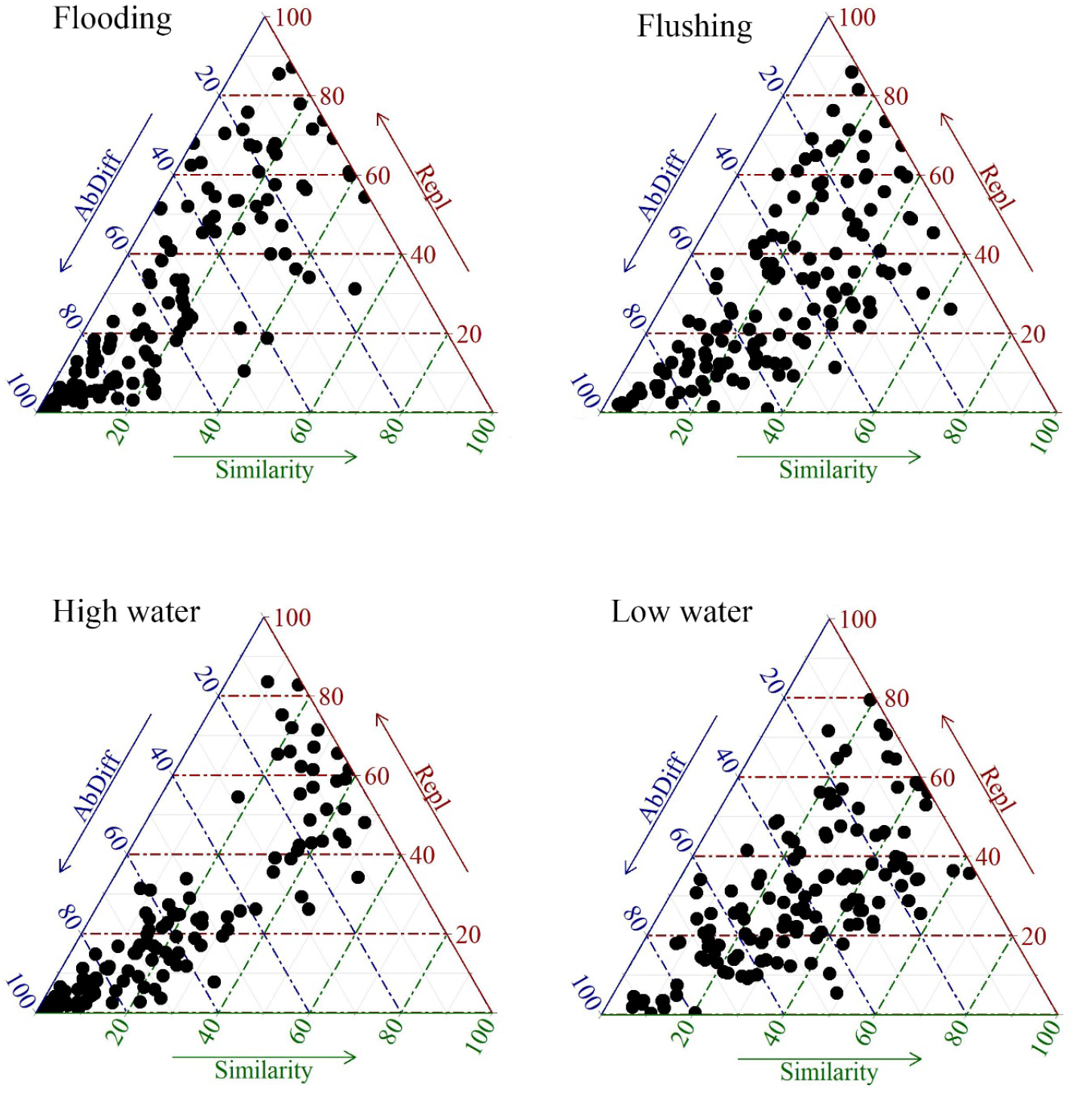
Triangular graph (simplex) of the proportion of elements in the beta diversity partition per pair of sample units for organism abundance values. AbDiff = abundance difference and Repl = species abundance replacement.

Because presented high collinearity or multicollinearity values, we removed the following environmental variables of each hydrological period: total chlorophyll, pH, conductivity and total dissolved solids (flooding); dissolved oxygen, pH, conductivity and total dissolved nitrogen (flushing); temperature, conductivity and total dissolved solids (high water) and dissolved oxygen, blue-green algae, pH, conductivity and total dissolved nitrogen (low water).

The environmental and spatial variables showed little influence on the distribution patterns of beta diversity and its components, regardless the hydrological period (Table 3). Considering the presence-absence species data, the environmental variables explained the beta diversity patterns in flushing and low waters periods and the richness difference component in in the low water period. Regarding the abundance data, the environmental variables explained the beta diversity and the abundance difference component in the high water period (Table 3).

**Table 3.**
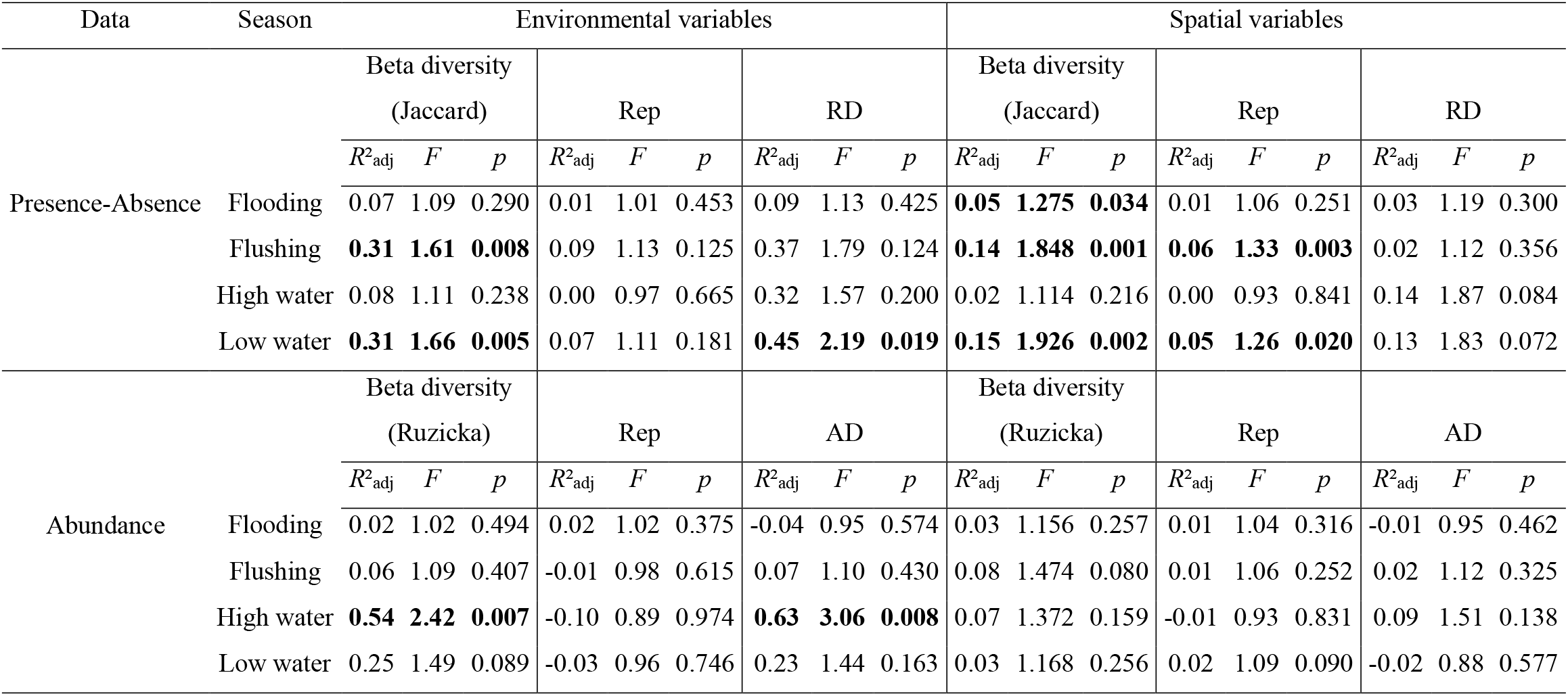
Distance-based redundancy analysis (dbRDA) of the influence of environmental and spatial predictors on the matrices resulting from the beta diversity partition. Rep = replacement; RD = richness difference; AD = abundance difference. Significant values are in bold

Concerning the presence-absence values, spatial variables explained the beta diversity patterns in flooding, flushing, and low water periods, and replacement component in flushing and low water periods. However, concerning the abundance data, spatial variables did not explain beta diversity nor its components in any of the hydrological periods analyzed (Table 3).

Regarding the concordance analyzes, zooplankton beta diversity and its components showed low values between hydrological periods (Table 4). Taking into account the presence and absence species data, there was concordance of beta diversity only in the comparisons between low water and flushing periods, and low water and high water periods (Table 4). Concerning the beta diversity components, there was concordance only in the comparisons between high water and low water (richness difference component) and between flooding and flushing (richness replacement component) and flooding and low water (richness replacement component). On the other hand, the abundance data did not show concordant patterns between the hydrological periods.

**Table 4.**
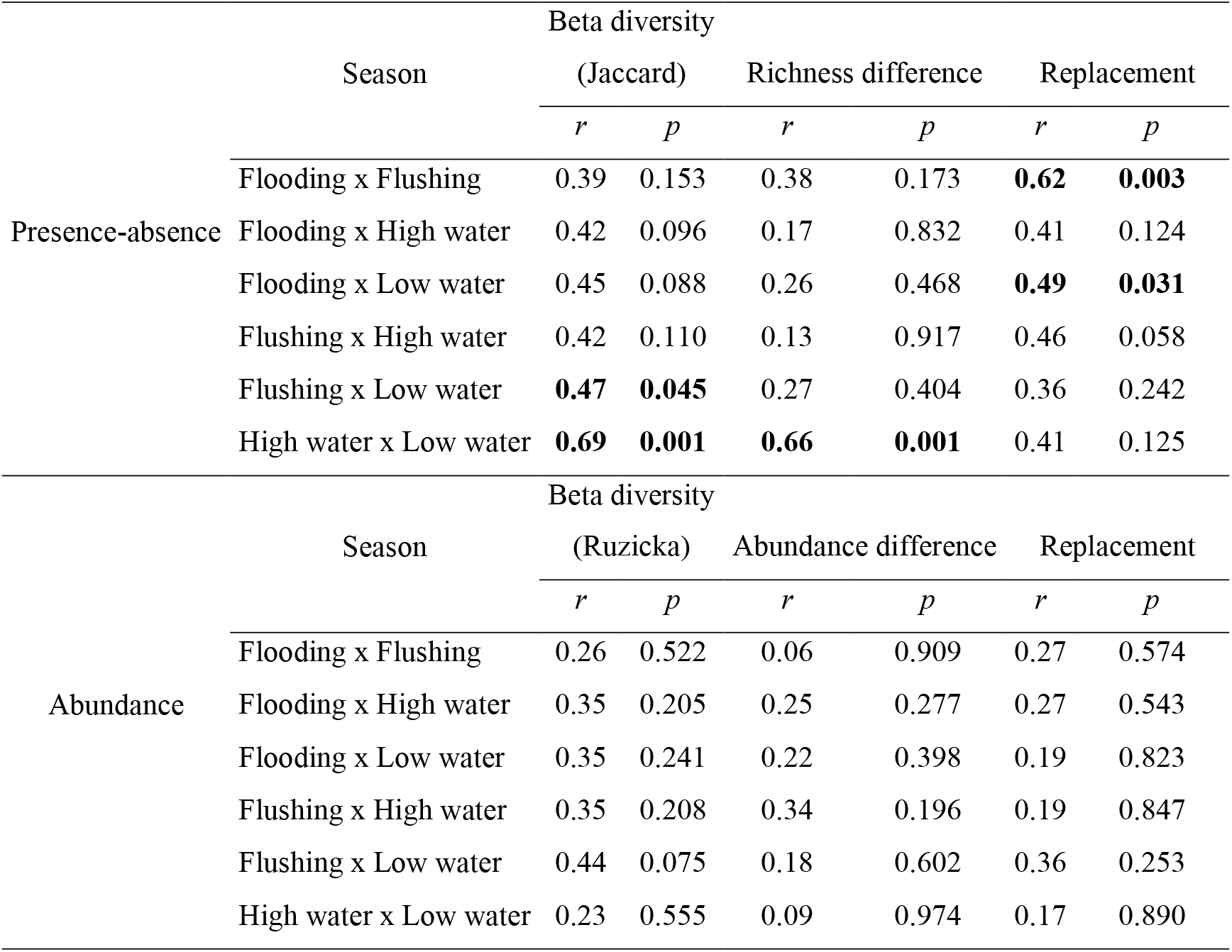
Procrustes test evaluating the concordance of beta diversity and its components values between hydrological periods. Significant values are in bold

|  |  | Beta diversity |  |  |  |  |  |
| --- | --- | --- | --- | --- | --- | --- | --- |
| Season |  | (Jaccard) |  | Richness difference |  | Replacement |  |
|  |  | *r* | *p* | *r* | *p* | *r* | *p* |
| Presence-absence | Flooding x Flushing | 0.39 | 0.153 | 0.38 | 0.173 | **0.62** | **0.003** |
|  | Flooding x High water | 0.42 | 0.096 | 0.17 | 0.832 | 0.41 | 0.124 |
|  | Flooding x Low water | 0.45 | 0.088 | 0.26 | 0.468 | **0.49** | **0.031** |
|  | Flushing x High water | 0.42 | 0.110 | 0.13 | 0.917 | 0.46 | 0.058 |
|  | Flushing x Low water | **0.47** | **0.045** | 0.27 | 0.404 | 0.36 | 0.242 |
|  | High water x Low water | **0.69** | **0.001** | **0.66** | **0.001** | 0.41 | 0.125 |
|  |  | Beta diversity |  |  |  |  |  |
| Season |  | (Ruzicka) |  | Abundance difference |  | Replacement |  |
|  |  | *r* | *p* | *r* | *p* | *r* | *p* |
| Abundance | Flooding x Flushing | 0.26 | 0.522 | 0.06 | 0.909 | 0.27 | 0.574 |
|  | Flooding x High water | 0.35 | 0.205 | 0.25 | 0.277 | 0.27 | 0.543 |
|  | Flooding x Low water | 0.35 | 0.241 | 0.22 | 0.398 | 0.19 | 0.823 |
|  | Flushing x High water | 0.35 | 0.208 | 0.34 | 0.196 | 0.19 | 0.847 |
|  | Flushing x Low water | 0.44 | 0.075 | 0.18 | 0.602 | 0.36 | 0.253 |
|  | High water x Low water | 0.23 | 0.555 | 0.09 | 0.974 | 0.17 | 0.890 |

## Discussion

### Local contributions to beta diversity

When evaluating the beta diversity local contributions patterns using the presence-absence data, we found that the main contribution sites were located in the south during flooding and low water periods. This lake region has a higher proportion of areas with pastoral use (Peres, Gurgel & Laques, 2018) and also the highest proportion of *igarapés* area. On the other hand, the northern region connects more predominantly with the Amazon River (Bonnet *et al*., 2008). Given that the variation in species composition and abundance influence the LCBD contributions, the land use may have influenced the difference in species composition between sites, what, consequently, influenced the increase the beta diversity contribution.

The significant LCBD site located in the southern region presented the lowest richness of individuals per sampling unit during the flooding period, while in the flushing period, it showed a different occurrence of species comparing to the other sites. High LCBD values may not be directly associated with high richness or abundance values, since areas with low richness and occurrences of differentiated species may also present higher contribution values, which may denote these areas as priorities for species conservation (Legendre & De Cáceres, 2013).

In the flooding and flushing periods, some sites in the northern region were differentiated concerning organism abundance. There was low abundance of zooplanktonic organisms at the significant site during the flooding period. Moreover, in flushing, the sampling unit ten, which most contributed to the beta diversity, stood out for the occurrence of different species compared to others sampling units in the same period (e.g., *Lecane elsa, Lecane luna* and *Nebela collaris*). In the low water period, the sampling unit also showed distinct species (e.g., *Difflugia elegans*). This distinction in the diversity patterns of the sampling units by hydrological period showed that the flood pulse promoted different dynamics in the floodplain lake. In the low water period, the sampling units were isolated from the main river, which means that the considerable environmental heterogeneity may have been influenced differences in species with different characteristics in each sampling period (Thomaz *et al*., 2007). In this case, as the sampling unit two, which has a higher LCBD value, is on the opposite side of the most important contribution area of the river’s water flow, located to the east, the isolation of the site may have justified such differentiation.

Whereas zooplanktonic organisms respond effectively to environmental variations (Vieira *et al*., 2011; Wang *et al*., 2016) and even greater impacts such as hydrological changes in cases of dams (Souza *et al*., 2019), we consider that the sampling units highlighted accordingly to the criteria of uniqueness by the LCBD analysis, being always associated with the marginal regions of the lake. These regions have higher interactions with *igarapés* and are in contact with the aquatic-terrestrial transition zones. Therefore, despite the hydrological importance of the flood pulse over the lake and the control over ecological dynamics, it is also important to take into account the importance that these *igarapés* and vegetation areas have for the existence of unique sites in relation to biodiversity for the Lago Grande do Curuai.

### Beta diversity partition

Related to the presence-absence data, there was a predominance of replacement concerning total beta diversity. It means that, despite a greater constancy in species richness per sampling unit, the species composition between pairs of units was different. Species are expected to show a substitution pattern over large environmental gradients, depending on other factors such as ecological tolerance of species (Legendre, 2014). Some studies report the sensitivity of organisms in the zooplankton community to environmental variations (Vieira *et al*., 2011), in some cases responding through changes in the trophic structure of the community (Ejsmont-Karabin *et al*., 2018) and changes in reproductive rates and species composition in the presence of other organisms (e.g., fish) (Feniova *et al*., 2019).

The high water period showed the highest beta diversity values and species replacement rate. It differed from our expectations, since we expected a greater environmental homogeneity and consequent biological homogeneity, reflecting a higher biological similarity between the sites due to the flood pulse in the high water period and due to the greater interconnectivity between habitats (Thomaz *et al*., 2007; Bozelli *et al*., 2015). Despite this, the increase in beta diversity values may have been attributable to a greater interaction area with the floodplain that began during the flooding period (Junk *et al*., 1989) and continued to settle during the high water period. This same pattern may have justified the lower beta diversity and replacement values in low water and flushing periods where, despite the isolation of habitats promoted by the reduction in the water volume, consequently minimized the interaction with the floodplain region and the main river.

On the other hand, although the beta diversity patterns using abundance data were the same for the presence-absence data with the highest values in the high water and flooding periods, the abundance difference component predominated over the replacement component. These values denote that, despite a greater tendency to replace species along the environmental gradient, these species had wide variations in abundance values. It highlighted the importance of understanding the zooplankton community abundance variations that, despite the ability to respond to environmental variations (e.g., variation in trophic status and phosphorus concentration in water), is often overlooked in some ecological studies (García-Chicote, Armengol & Rojo, 2018).

### Environmental and spatial predictors

Despite the distinctions observed in the patterns of similarity and substitution of species between hydrological periods, we observed that the environmental variables showed little prediction about the diversity patterns of the zooplankton community for presence and absence data. These variables explained only the patterns of similarity in the flushing and low water periods and the richness difference in the low water period.

On the other hand, there was a higher pattern of prediction of spatial variables over patterns of similarity in the composition of species, not explaining only in the high water period. These values denote that spatial variation may have a greater control over the organisms composition dynamics than environmental variation. Despite this, this control was only related to presence-absence values. The patterns of organism abundance and presence-absence refer to different factors. For example, for presence and absence data, beta diversity corresponding to the inverse of similarity in the composition is prioritized (Podani & Schmera, 2011), while for abundance data, besides the composition, variations in the number of individuals of each species are also considered. Therefore, when abundance is taken into account, sites with high species dissimilarity values are those that present a high distinction in species composition and the corresponding organisms abundance (Podani *et al*., 2013).

Therefore, the explanation obtained in the low water period using the presence-absence data may be related to the heterogeneity of ecological niches (Legendre, 2014). The low water period may have promoted the existence of different niches, some with more species and others with fewer species, due to the isolation. The substitution of species explained spatially may also be based on the isolation that makes the species of an environment unable to reach other places (Thomaz *et al*., 2007). For this reason, spatial isolation can drive the pattern of differentiation of species within the habitat and this same pattern may explain the spatial prediction in the period of flushing.

For the beta diversity components using abundance data, there was a low standard of explanation for both environmental and spatial variables, which showed that there was a greater complexity of factors (e.g., competition and predation) that may have been the most responsible for these variations and that were not evaluated in this study. This low pattern of response shows that the zooplankton community is not responding only to environmental variations at that time, but to changes that occurred in other periods before the sampling carried out. Besides, as the abundance and presence-absence data responded differently to different factors, we emphasize that both approaches can be complementary when used for biological monitoring purposes.

### Temporal concordance between beta diversity components

Despite the occurrence of significant values when evaluating the temporal concordance between the beta diversity patterns using presence-absence values, no pair of periods showed concordance between all the diversity patterns over the hydrological cycle. There was also no concordance between the periods using the abundance values. These results are in agreement with our expectations since even in other environments, there is a low standard of predictability and synchrony of zooplankton with other variables that allow us to predict a constant and predictable pattern for this community (Vieira *et al*., 2019).

These results also show that the environmental and biological dynamics of the floodplains are complex to be predictable and, depending on the hydrological period, which changes the entrance of river sediments and the inflow or outflow of water in the floodplain, and the evaluated group, the structuring of the communities can be different (Amoros & Bornette, 2002). There are proposals that the dynamics are so distinct and susceptible to hydrological variations that the high water period acts as a resumption of the successional regime of the structure and composition of the zooplankton community (Baranyi *et al*., 2002; Bozelli *et al*., 2015). Therefore, despite the economic advantages of sampling in less hydrological periods, we found that, in order to understand the beta diversity patterns of the zooplankton community, sampling are necessary to occur in all the hydrological periods of high and low waters, as well as in the flood and ebb intermediate periods.

## Conclusions

Hydrological variations govern the zooplankton community dynamics, thus the contribution of different locations depending on the hydrological period evaluated. With some exceptions, the sites that most contributed to the beta diversity presented less organism richness or abundance and also showed proximity to the coastal regions of the lake, especially those associated with *Igarapés* when using the organisms presence-absence data. This result denotes the relevance of these areas for biological monitoring and for the delimitation of priority areas for the conservation of zooplankton diversity.

Beta diversity was greatest in flooding and high water periods. Despite the differences in the partition values by hydrological period, the species replacement was dominant in all hydrological periods using the organisms presence-absence data, while the abundance difference was dominant using the quantitative values of organisms per sample unit. Therefore, the studies must evaluate both abundance and presence-absence data as a complementary way, considering that they can portray different processes in the face of environmental and spatial variations. Due to the complexity of factors that govern the distribution of zooplankton organisms in floodplains, there was a little prediction of environmental and spatial variables on the beta diversity distribution patterns for the community. Also, there was a low concordance between the patterns for the different hydrological periods, which highlights the need to study the hydrological periods of high and low waters, as well as the transient periods of flooding and flushing to obtain an adequate assessment of the dynamics distribution patterns of the zooplankton community from the perspective of beta diversity.

## Acknowledgments

The authors thank the Coordenação de Aperfeiçoamento de Pessoal de Nível Superior (CAPES) for providing financial assistance to LFG, ACAMG, CAS and HRP. LCGV was supported by productivity fellowships of Conselho Nacional de Desenvolvimento Científico e Tecnológico (CNPq). We also thank the Fondation pour la Recherche sur la Biodiversité (FRB) and the Conselho Nacional de Desenvolvimento Científico e Tecnológico (CNPq); these groups, in partnership with Institut de Recherche pour le Développement (IRD), financed the project number: 490634/2013-3.

## Data availability statement

Data are available on request from the corresponding author.

## References

Affonso A., Barbosa C. & Novo E. (2011). Water quality changes in floodplain lakes due to the Amazon River flood pulse: Lago Grande de Curuaí (Pará). Brazilian Journal of Biology 71, 601–610. https://doi.org/10.1590/S1519-69842011000400004

Amoros C. & Bornette G. (2002). Connectivity and biocomplexity in waterbodies of riverine floodplains. Freshwater Biology 47, 761–776. https://doi.org/10.1046/j.1365-2427.2002.00905.x

APHA (2005). Standard Methods for the Examination of Water and Wastewater, 21st edn. American Public Health Association/American Water Works Association/Water Environment Federation, Washington DC.

Baranyi C., Hein T., Holarek C., Keckeis S. & Schiemer F. (2002). Zooplankton biomass and community structure in a Danube River floodplain system: effects of hydrology. Freshwater Biology 47, 473–482. https://doi.org/10.1046/j.1365-2427.2002.00822.x

Baselga A. (2010). Partitioning the turnover and nestedness components of beta diversity. Global Ecology and Biogeography 19, 134–143. https://doi.org/10.1111/j.1466-8238.2009.00490.x

Bonnet M.P., Barroux G., Martinez J.M., Seyler F., Moreira-Turcq P., Cochonneau G., et al. (2008). Floodplain hydrology in an Amazon floodplain lake (Lago Grande de Curuaí). Journal of Hydrology 349, 18–30. https://doi.org/10.1016/j.jhydrol.2007.10.055

Borcard D., Gillet F. & Legendre P. (2018). Numerical Ecology with R. Springer International Publishing, Cham.

Bottrell H.H., Duncan A., Gliwicz Z.M., Grygierek E., Herzig A., Hillbrichtilkowska A., et al. (1976). Review of some problems in zooplankton production studies. Norwegian Journal of Zoology 24, 419–456

Bozelli R.L., Thomaz S.M., Padial A.A., Lopes P.M. & Bini L.M. (2015). Floods decrease zooplankton beta diversity and environmental heterogeneity in an Amazonian floodplain system. Hydrobiologia 753, 233–241. https://doi.org/10.1007/s10750-015-2209-1

Carneiro F.M., Nabout J.C., Vieira L.C.G., Lodi S. & Bini L.M. (2013). Higher Taxa Predict Plankton Beta-diversity Patterns Across an Eutrophication Gradient. Natureza & Conservação 11, 43–47. https://doi.org/10.4322/natcon.2013.006

Chambers J.M. (2013). SoDA: Functions and Examples for “Software for Data Analysis”

Dray S., Blanchet G., Borcard D., Clappe S., Guenard G., Jombart T., et al. (2018). adespatial: Multivariate Multiscale Spatial Analysis. R package version 0.1-1

Dray S., Legendre P. & Peres-Neto P.R. (2006). Spatial modelling: a comprehensive framework for principal coordinate analysis of neighbour matrices (PCNM). Ecological Modelling 196, 483–493. https://doi.org/10.1016/j.ecolmodel.2006.02.015

Ejsmont-Karabin J., Feniova I., Kostrzewska-Szlakowska I., Rzepecki M., Petrosyan V.G. & Dzialowski A.R. (2018). Factors influencing phosphorus regeneration by lake zooplankton—An experimental approach. Limnologica 70, 58–64. https://doi.org/10.1016/j.limno.2018.01.003

Feniova I., Sakharova E., Karpowicz M., Gladyshev M.I., Sushchik N.N., Dawidowicz P., et al. (2019). Direct and Indirect Impacts of Fish on Crustacean Zooplankton in Experimental Mesocosms. Water 11, 2090. https://doi.org/10.3390/w11102090

García-Chicote J., Armengol X. & Rojo C. (2018). Zooplankton abundance: A neglected key element in the evaluation of reservoir water quality. Limnologica 69, 46–54. https://doi.org/10.1016/j.limno.2017.11.004

Gower J.C. (1975). Generalized procrustes analysis. Psychometrika 40, 33–51. https://doi.org/10.1007/BF02291478

Guisan A. & Thuiller W. (2005). Predicting species distribution: offering more than simple habitat models. Ecology Letters 8, 993–1009. https://doi.org/10.1111/j.1461-0248.2005.00792.x

Junk W.J., Bayley P.B. & Sparks R.E. (1989). The flood pulse concept in river-floodplain systems. Canadian special publication of fisheries and aquatic sciences 106, 110–127

Junk W.J., Piedade M.T.F., Schöngart J., Cohn-Haft M., Adeney J.M. & Wittmann F. (2011). A Classification of Major Naturally-Occurring Amazonian Lowland Wetlands. Wetlands 31, 623–640. https://doi.org/10.1007/s13157-011-0190-7

Junk W.J., Piedade M.T.F., Schöngart J. & Wittmann F. (2012). A classification of major natural habitats of Amazonian white-water river floodplains (várzeas). Wetlands Ecology and Management 20, 461–475. https://doi.org/10.1007/s11273-012-9268-0

Legendre P. (2014). Interpreting the replacement and richness difference components of beta diversity. Global Ecology and Biogeography 23, 1324–1334. https://doi.org/10.1111/geb.12207

Legendre P. & Andersson M.J. (1999). Distance-based redundancy analysis: Testing multispecies responses in multifactorial ecological experiments. Ecological Monographs 69, 1–24. https://doi.org/10.1890/0012-9615(1999)069[0001:DBRATM]2.0.CO;2

Legendre P. & De Cáceres M. (2013). Beta diversity as the variance of community data: dissimilarity coefficients and partitioning. Ecology Letters 16, 951–963. https://doi.org/10.1111/ele.12141

Legendre P. & Legendre L. (2012). Numerical ecology, Third. Elsevier, Amsterdam.

de Moraes Novo E.M.L., de Farias Barbosa C.C., de Freitas R.M., Shimabukuro Y.E., Melack J.M. & Filho W.P. (2006). Seasonal changes in chlorophyll distributions in Amazon floodplain lakes derived from MODIS images. Limnology 7, 153–161. https://doi.org/10.1007/s10201-006-0179-8

de Morais G.F., dos Santos Ribas L.G., Ortega J.C.G., Heino J. & Bini L.M. (2018). Biological surrogates: A word of caution. Ecological Indicators 88, 214–218. https://doi.org/10.1016/j.ecolind.2018.01.027

Oksanen J., Blanchet, Guillaume F., Friendly M., Kindt R., Legendre P., Mcglinn D., et al. (2016). Vegan: Community Ecology Package. R package version 2.4-0

Oksanen J., Blanchet F.G., Kindt R., Legendre P., Minchin P.R., O’hara R.B., et al. (2013). Package ‘vegan.’ Community ecology package, version 2

Peres L.G.M., Gurgel H. & Laques A.-E. (2018). Dinâmica da paisagem em planícies de inundação amazônicas: o caso do Lago Grande do Curuai, Pará, Brasil. Confins. Revue franco-brésilienne de géographie/Revista franco-brasilera de geografia

Podani J., Ricotta C. & Schmera D. (2013). A general framework for analyzing beta diversity, nestedness and related community-level phenomena based on abundance data. Ecological Complexity 15, 52–61. https://doi.org/10.1016/j.ecocom.2013.03.002

Podani J. & Schmera D. (2011). A new conceptual and methodological framework for exploring and explaining pattern in presence - absence data. Oikos 120, 1625–1638. https://doi.org/10.1111/j.1600-0706.2011.19451.x

R Core Team (2016). R: A Language and Environment for Statistical Computing

Socolar J.B., Gilroy J.J., Kunin W.E. & Edwards D.P. (2016). How Should Beta-Diversity Inform Biodiversity Conservation? Trends in Ecology & Evolution 31, 67–80. https://doi.org/10.1016/j.tree.2015.11.005

Souza C.A. de, Vieira L.C.G., Legendre P., Carvalho P. de, Velho L.F.M. & Beisner B.E. (2019). Damming interacts with the flood pulse to alter zooplankton communities in an Amazonian river. Freshwater Biology, fwb.13284. https://doi.org/10.1111/fwb.13284

Thomaz S.M., Bini L.M. & Bozelli R.L. (2007). Floods increase similarity among aquatic habitats in river-floodplain systems. Hydrobiologia 579, 1–13. https://doi.org/10.1007/s10750-006-0285-y

Vieira A.C.B., Medeiros A.M.A., Ribeiro L.L. & Crispim M.C. (2011). Population dynamics of Moina minuta Hansen (1899), Ceriodaphnia cornuta Sars (1886), and Diaphanosoma spinulosum Herbst (1967) (Crustacea: Branchiopoda) in different nutrients (N and P) concentration ranges. Acta Limnologica Brasiliensia 23, 48–56. https://doi.org/10.4322/actalb.2011.018

Vieira M.C., Bini L.M., Velho L.F.M., Gomes L.F., Nabout J.C. & Vieira L.C.G. (2017). Biodiversity shortcuts in biomonitoring of novel ecosystems. Ecological Indicators 82, 505–512. https://doi.org/10.1016/j.ecolind.2017.07.025

Vieira M.C., Roitman I., de Oliveira Barbosa H., Machado Velho L.F. & Cardoso Galli Vieira L. (2019). Spatial synchrony of zooplankton during the impoundment of amazonic reservoir. Ecological Indicators 98, 649–656. https://doi.org/10.1016/j.ecolind.2018.11.040

Wang C., Wang L., Deng D. & Zhou Z. (2016). Temporal and spatial variations in rotifer correlations with environmental factors in Shengjin Lake, China. Environmental Science and Pollution Research 23, 8076–8084. https://doi.org/10.1007/s11356-015-6009-y

Whittaker R.H. (1960). Vegetation of the Siskiyou Mountains, Oregon and California. Ecological Monographs 30, 279–338. https://doi.org/10.2307/1943563

