## Supplementary material for "Zooplankton community beta diversity in an Amazonian floodplain lake"

**Table S1** – Mean density of zooplankton taxa identified for each sampling period. SD= Standard Deviation.

| Group | Specie | Flooding | | Flushing | | High water | | Low water | |
|---|---|---|---|---|---|---|---|---|---|
| | | Mean | SD | Mean | SD | Mean | SD | Mean | SD |
| Cladocera | *Acroperus tupinamba Sinev & Elmoor-Loureiro, 2010* | 0.0 | 0.0 | 0.0 | 0.0 | 0.9 | 3.6 | 0.0 | 0.0 |
| | *Alona guttata* Sars, 1862 | 9.4 | 36.3 | 14.7 | 60.6 | 18.9 | 32.9 | 312.0 | 682.7 |
| | *Alona ossiani* Sinev, 1998 | 0.0 | 0.0 | 0.0 | 0.0 | 2.5 | 10.1 | 0.0 | 0.0 |
| | *Alonella dadayi* Birge, 1910 | 0.0 | 0.0 | 19.6 | 80.8 | 0.0 | 0.0 | 0.0 | 0.0 |
| | *Anthalona verrucosa* Sars, 1901 | 0.0 | 0.0 | 0.0 | 0.0 | 6.0 | 20.3 | 0.0 | 0.0 |
| | *Bosmina hagmanni* Stingelin, 1904 | 262.5 | 405.4 | 676.7 | 1205.8 | 166.9 | 299.5 | 1623.7 | 3825.3 |
| | *Bosmina tubicen* Brehm, 1953 | 160.7 | 275.2 | 649.6 | 933.7 | 95.5 | 272.9 | 64.1 | 142.7 |
| | *Bosminopsis deitersi* Richard, 1895 | 1107.4 | 2110.3 | 5809.8 | 14372.2 | 883.2 | 1433.8 | 0.0 | 0.0 |
| | *Ceriodaphnia cornuta* Sars, 1885 | 1882.5 | 2449.9 | 457.1 | 606.5 | 555.7 | 1541.0 | 923.5 | 2375.4 |
| | *Ceriodaphnia reticulata* Jurine, 1820 | 76.0 | 150.4 | 23.5 | 97.0 | 0.0 | 0.0 | 0.0 | 0.0 |
| | *Ceriodaphnia silvestrii* Daday, 1902 | 88.6 | 363.7 | 14.7 | 60.6 | 0.0 | 0.0 | 0.0 | 0.0 |
| | *Chydorus eurynotus* Sars, 1901 | 0.4 | 1.1 | 0.0 | 0.0 | 12.5 | 24.2 | 0.0 | 0.0 |
| | *Chydorus pubescens* Sars, 1901 | 117.6 | 485.1 | 0.0 | 0.0 | 0.0 | 0.0 | 0.0 | 0.0 |
| | *Chydorus sphaericus* Elmoor-Loureiro, 1997 | 7.4 | 30.3 | 0.0 | 0.0 | 0.0 | 0.0 | 0.0 | 0.0 |
| | *Coronatella monacantha* Sars, 1901 | 0.0 | 0.0 | 0.0 | 0.0 | 3.7 | 15.2 | 0.4 | 1.6 |
| | *Coronatella poppei* Richard, 1897 | 0.0 | 0.0 | 0.0 | 0.0 | 0.0 | 0.0 | 0.2 | 0.8 |
| | *Diaphanosoma birgei* Korinek, 1981 | 1690.5 | 2721.2 | 254.9 | 487.1 | 59.3 | 111.6 | 2118.0 | 3533.2 |
| | *Diaphanosoma spinulosum* Herbst, 1975 | 1067.9 | 1367.8 | 16.1 | 60.3 | 261.4 | 603.4 | 294.1 | 1212.7 |
| | *Grimaldina brazzai* Richard, 1892 | 0.0 | 0.0 | 0.0 | 0.0 | 1.2 | 5.1 | 0.0 | 0.0 |
| | *Holopedium amazonicum* Stingelin, 1904 | 1552.2 | 3165.5 | 0.0 | 0.0 | 0.0 | 0.0 | 0.0 | 0.0 |
| | *Karualona muelleri* Richard, 1897 | 0.0 | 0.0 | 25.5 | 105.1 | 0.0 | 0.0 | 20.6 | 80.6 |
| | *Leydigiopsis megalops* Sars, 1901 | 0.0 | 0.0 | 0.0 | 0.0 | 0.2 | 0.8 | 0.0 | 0.0 |
| | *Macrothrix laticornis* Jurine, 1820 | 0.2 | 0.8 | 25.5 | 105.1 | 7.8 | 32.3 | 40.4 | 161.4 |
| | *Macrothrix mira* Smirnov, 1992 | 14.7 | 60.6 | 0.0 | 0.0 | 0.0 | 0.0 | 0.4 | 1.6 |
| | *Moina micrura* Kurz, 1875 | 274.9 | 519.7 | 1250.0 | 2318.0 | 163.5 | 565.3 | 344.1 | 707.8 |

| | | | | | | | | | |
|---|---|---|---|---|---|---|---|---|---|
| | *Moina minuta* Hansen, 1899 | 1580.6 | 4727.4 | 701.4 | 1128.3 | 92.1 | 238.6 | 3911.2 | 4176.3 |
| | *Moina reticulata* Daday, 19*05* | 0.0 | 0.0 | 23.5 | 97.0 | 0.0 | 0.0 | 0.0 | 0.0 |
| | *Nicsmirnovius incredibilis* Smirnov, 1984 | 0.0 | 0.0 | 0.0 | 0.0 | 3.5 | 14.6 | 0.0 | 0.0 |
| | *Picripleuroxus similis* Vávra, 1900 | 3.9 | 15.1 | 0.0 | 0.0 | 0.0 | 0.0 | 0.0 | 0.0 |
| | *Simocephalus* sp. Orlova-Bienkowskaja, 2001 | 0.0 | 0.0 | 0.0 | 0.0 | 0.0 | 0.0 | 24.5 | 101.1 |
| | *Streblocerus pygmaeus* Sars, 190*1* | 0.0 | 0.0 | 0.0 | 0.0 | 0.2 | 0.8 | 0.0 | 0.0 |
| Copepoda | *Argyrodiaptomus azevedoi* Wright, 1935 | 44.1 | 132.1 | 0.0 | 0.0 | 0.0 | 0.0 | 0.0 | 0.0 |
| | *Argyrodiaptomus robertsonae* Dussart, 1985 | 832.2 | 3236.0 | 0.0 | 0.0 | 74.5 | 307.2 | 0.0 | 0.0 |
| | Cyclopidae copepodit | 6352.8 | 6972.1 | 8217.6 | 6887.5 | 6274.4 | 13228.9 | 6064.7 | 4475.1 |
| | Cyclopidae *nauplii* | 10291.8 | 9572.5 | 53820.6 | 40730.0 | 8928.7 | 14298.1 | 24213.7 | 15382.4 |
| | Diaptomidae copepodit | 6575.5 | 11675.1 | 1245.1 | 1457.9 | 1194.3 | 2790.3 | 3725.5 | 2975.0 |
| | Diaptomidae *nauplii* | 14521.1 | 29058.7 | 1904.0 | 1777.7 | 934.6 | 2617.8 | 3444.1 | 3265.9 |
| | *Diaptomus deitersi Poppe, 1891* | 6.4 | 26.5 | 0.0 | 0.0 | 0.4 | 1.6 | 0.0 | 0.0 |
| | *Mesocyclops meridianus Kiefer, 1926* | 10.3 | 31.9 | 0.0 | 0.0 | 0.0 | 0.0 | 0.0 | 0.0 |
| | *Metacyclops mendocinus* Wierzejski, 1892 | 14.7 | 60.6 | 0.0 | 0.0 | 39.6 | 161.6 | 19.6 | 80.8 |
| | *Microcyclops alius* Kiefer, 1935 | 116.2 | 388.0 | 26.9 | 110.9 | 826.2 | 3309.5 | 470.8 | 866.4 |
| | *Microcyclops anceps* Richard, 189*7* | 0.0 | 0.0 | 0.0 | 0.0 | 14.1 | 27.1 | 259.8 | 440.8 |
| | *Microcyclops ceibaensis* Marsh, 1919 | 0.0 | 0.0 | 138.2 | 457.2 | 93.1 | 384.0 | 0.0 | 0.0 |
| | *Microcyclops finitimus* Dussart, 1984 | 232.4 | 522.6 | 14.7 | 60.6 | 0.0 | 0.0 | 0.0 | 0.0 |
| | *Microcyclops* sp. Claus, 1893 | 0.0 | 0.0 | 25.7 | 105.1 | 17.6 | 64.7 | 47.1 | 194.0 |
| | *Notodiaptomus amazonicus* Wright, 1935 | 115.6 | 246.1 | 40.2 | 118.0 | 0.0 | 0.0 | 731.3 | 882.2 |
| | *Notodiaptomus kieferi* Brandorff, 1973 | 29.4 | 121.3 | 0.0 | 0.0 | 0.0 | 0.0 | 0.0 | 0.0 |
| | *Notodiaptomus paraensis* Dussart & Robertson B.A., 1984 | 22.7 | 67.1 | 0.0 | 0.0 | 23.0 | 77.8 | 0.0 | 0.0 |
| | *Thermocyclops decipiens* Kiefer, 1929 | 0.0 | 0.0 | 2330.2 | 3146.4 | 470.9 | 1918.8 | 2665.1 | 2149.4 |
| | *Thermocyclops inversus* Kiefer, 1936 | 155.9 | 416.4 | 0.2 | 0.8 | 354.9 | 1454.7 | 431.4 | 537.0 |
| | *Thermocyclops minutus* Lowndes, 1934 | 0.0 | 0.0 | 0.0 | 0.0 | 7.8 | 32.3 | 39.2 | 161.7 |
| | *Thermocyclops* sp. Kiefer, 1927 | 0.0 | 0.0 | 0.0 | 0.0 | 48.5 | 163.8 | 0.0 | 0.0 |

| | | | | | | | | | |
|---|---|---|---|---|---|---|---|---|---|
| Rotifera | *Ascomorpha agilis* Zacharias, 1893 | 0.0 | 0.0 | 0.0 | 0.0 | 49.0 | 202.1 | 24.5 | 101.1 |
| | *Ascomorpha eucadis* Perty, 1850 | 225.5 | 635.2 | 0.0 | 0.0 | 0.0 | 0.0 | 706.3 | 1247.7 |
| | *Ascomorpha saltans* Bartsch, 1870 | 39.2 | 161.7 | 0.0 | 0.0 | 308.9 | 664.6 | 455.9 | 912.0 |
| | *Ascomorpha* sp. Perty, 1850 | 0.0 | 0.0 | 0.0 | 0.0 | 0.0 | 0.0 | 235.3 | 482.5 |
| | *Asplanchna herricki* Guerne, 1888 | 0.0 | 0.0 | 0.0 | 0.0 | 0.0 | 0.0 | 156.9 | 646.8 |
| | *Asplanchna priodonta* Gosse, 1850 | 0.0 | 0.0 | 0.0 | 0.0 | 0.0 | 0.0 | 98.0 | 404.2 |
| | *Asplanchna sieboldii* Leydig, 1854 | 3.7 | 15.2 | 1105.9 | 2855.7 | 1.2 | 2.9 | 1001.6 | 4037.6 |
| | *Asplanchna* sp. Gosse, 1850 | 0.0 | 0.0 | 0.0 | 0.0 | 0.0 | 0.0 | 19.8 | 80.8 |
| | *Bdelloidea* Hudson, 1884 | 0.0 | 0.0 | 535.3 | 2098.0 | 0.0 | 0.0 | 24.5 | 101.1 |
| | *Beauchampiella eudactylota* Gosse, 1886 | 0.0 | 0.0 | 0.0 | 0.0 | 5.8 | 14.0 | 0.0 | 0.0 |
| | *Brachionus ahlstromi Lindeman, 1939* | 14.7 | 60.6 | 338.2 | 1394.6 | 0.0 | 0.0 | 0.0 | 0.0 |
| | *Brachionus bidentatus* Anderson, 1889 | 0.0 | 0.0 | 25.5 | 105.1 | 0.0 | 0.0 | 29.4 | 121.3 |
| | *Brachionus calyciflorus* Pallas, 1766 | 1564.7 | 4927.6 | 5646.5 | 12541.4 | 0.6 | 1.8 | 1764.9 | 2572.3 |
| | *Brachionus caudatus* Barrois & Daday, 1894 | 0.0 | 0.0 | 6629.4 | 11937.7 | 0.0 | 0.0 | 12685.5 | 18252.8 |
| | *Brachionus dolabratus* Harring, 1914 | 7.5 | 30.3 | 4300.6 | 8131.4 | 41.4 | 110.2 | 39.4 | 161.6 |
| | *Brachionus falcatus* Zacharias, 1898 | 227.3 | 455.5 | 784.3 | 1340.5 | 79.0 | 323.2 | 153.1 | 340.8 |
| | *Brachionus mirus* Daday, 1905 | 7.4 | 30.3 | 744.1 | 1369.3 | 19.6 | 80.8 | 753.5 | 1525.0 |
| | *Brachionus urceolaris* Müller, 1773 | 0.0 | 0.0 | 0.0 | 0.0 | 0.0 | 0.0 | 19.6 | 80.8 |
| | *Brachionus zahniseri* Ahlstrom, 1934 | 0.0 | 0.0 | 15703.9 | 27220.6 | 408.4 | 1209.1 | 78.6 | 323.3 |
| | *Cephalodella* cf. *catellina* Müller, 1786 | 0.0 | 0.0 | 0.0 | 0.0 | 0.0 | 0.0 | 1294.7 | 2504.9 |
| | *Cephalodella hoodii* Gosse, 1886 | 0.0 | 0.0 | 29.4 | 121.3 | 0.0 | 0.0 | 0.0 | 0.0 |
| | *Cephalodella* sp. Bory de St. Vincent, 1826 | 0.0 | 0.0 | 0.0 | 0.0 | 0.0 | 0.0 | 117.6 | 332.1 |
| | *Cephalodella tenuiseta Burn, 1890* | 4.9 | 20.2 | 0.0 | 0.0 | 0.0 | 0.0 | 0.0 | 0.0 |
| | *Collotheca edentata* Collins, 1872 | 3.7 | 15.2 | 24.5 | 101.1 | 0.0 | 0.0 | 0.0 | 0.0 |
| | *Collotheca mutabilis* Hudson, 1885 | 29.4 | 121.3 | 0.0 | 0.0 | 0.0 | 0.0 | 0.0 | 0.0 |
| | *Collotheca stephanochaeta* Edmondson, 1936 | 22.1 | 91.0 | 0.0 | 0.0 | 0.0 | 0.0 | 0.0 | 0.0 |
| | *Collotheca* sp. Harring, 1913 | 0.0 | 0.0 | 0.0 | 0.0 | 3.9 | 16.2 | 0.0 | 0.0 |

| | | | | | | | | |
|---|---|---|---|---|---|---|---|---|
| *Collotheca tubiformis* Nipkow, 1961 | 29.4 | 121.3 | 0.0 | 0.0 | 0.0 | 0.0 | 0.0 | 0.0 |
| *Collotheca undulata* Sládeček, 1969 | 0.0 | 0.0 | 0.0 | 0.0 | 19.6 | 80.8 | 0.0 | 0.0 |
| *Colurella Hindenburgi* Steinecke, 1916 | 0.0 | 0.0 | 19.6 | 80.8 | 0.0 | 0.0 | 0.0 | 0.0 |
| *Colurella obtusa* Gosse, 1886 | 0.0 | 0.0 | 994.5 | 3014.7 | 0.0 | 0.0 | 0.0 | 0.0 |
| *Colurella* sp. Bory de St. Vincent, 1824 | 0.0 | 0.0 | 19.6 | 80.8 | 0.0 | 0.0 | 0.0 | 0.0 |
| *Conochilus* sp. Ehrenberg, 1834 | 0.0 | 0.0 | 0.0 | 0.0 | 0.0 | 0.0 | 19.6 | 80.8 |
| *Conochilus unicornis* Rousselet, 1892 | 3251.6 | 7934.5 | 1.9 | 5.5 | 1039.2 | 3655.1 | 0.0 | 0.0 |
| *Dicranophorus forcipatus* Müller, 1786 | 0.0 | 0.0 | 0.0 | 0.0 | 13.2 | 33.4 | 0.0 | 0.0 |
| *Dicranophorus* sp. O.F. Muller, 1773 | 0.0 | 0.0 | 14.7 | 60.6 | 0.0 | 0.0 | 0.0 | 0.0 |
| *Drilophaga delagei* Beauchamp, 1904 | 253.7 | 769.4 | 0.0 | 0.0 | 0.0 | 0.0 | 0.0 | 0.0 |
| *Elosa worrali* Lord, 1891 | 0.0 | 0.0 | 0.0 | 0.0 | 152.3 | 466.2 | 0.0 | 0.0 |
| *Epiphanes clavatula* Ehrenberg, 1832 | 411.8 | 1697.7 | 984.3 | 2779.0 | 0.0 | 0.0 | 0.0 | 0.0 |
| *Epiphanes macrourus* Barrois & Daday, 1894 | 400.5 | 1029.6 | 0.0 | 0.0 | 0.0 | 0.0 | 3681.8 | 11172.4 |
| *Epiphanes pelagica* Jennings, 1900 | 2.5 | 10.1 | 0.0 | 0.0 | 0.0 | 0.0 | 0.0 | 0.0 |
| *Euchlanis meneta* Myers, 1930 | 0.0 | 0.0 | 0.0 | 0.0 | 2.5 | 10.1 | 0.0 | 0.0 |
| *Euchlanis* sp. Ehrenberg, 1832 | 0.0 | 0.0 | 0.0 | 0.0 | 0.0 | 0.0 | 78.4 | 323.4 |
| *Euchlanis triquetra* Ehrenberg, 1838 | 0.0 | 0.0 | 0.0 | 0.0 | 123.5 | 267.7 | 156.9 | 501.6 |
| *Filinia camasecla* Myers, 1938 | 0.0 | 0.0 | 278.8 | 441.1 | 2.0 | 4.4 | 0.0 | 0.0 |
| *Filinia longiseta* Ehrenberg, 1834 | 101.9 | 261.2 | 4502.0 | 4730.1 | 33.0 | 85.7 | 7698.4 | 10641.8 |
| *Filinia opoliensis* Zacharias, 1898 | 14.7 | 60.6 | 100.0 | 308.2 | 40.0 | 161.5 | 39.6 | 161.6 |
| *Filinia terminalis* Plate, 1886 | 105.6 | 200.9 | 102.0 | 420.4 | 21.7 | 76.5 | 59.0 | 242.5 |
| *Filinia* sp Bory de St. Vincent, 1824 | 14.7 | 60.6 | 0.0 | 0.0 | 0.0 | 0.0 | 0.0 | 0.0 |
| *Gastropus hyptopus* Ehrenberg, 1838 | 0.0 | 0.0 | 485.3 | 1937.4 | 121.6 | 423.1 | 0.0 | 0.0 |
| *Gastropus stylifer* Imhof, 1891 | 0.0 | 0.0 | 0.0 | 0.0 | 1.6 | 6.5 | 0.0 | 0.0 |
| *Harringia eupoda* Gosse, 1887 | 897.1 | 3698.7 | 0.0 | 0.0 | 0.0 | 0.0 | 0.0 | 0.0 |
| *Hexarthra intermedia* Wiszniewski, 1929 | 0.0 | 0.0 | 320.6 | 1056.1 | 0.0 | 0.0 | 0.0 | 0.0 |
| *Hexarthra* cf. *mira* Hudson, 1871 | 0.0 | 0.0 | 117.6 | 376.2 | 0.0 | 0.0 | 0.0 | 0.0 |

| | | | | | | | | |
|---|---|---|---|---|---|---|---|---|
| *Hexarthra* sp. Schmarda, 1854 | 0.0 | 0.0 | 235.3 | 970.1 | 0.0 | 0.0 | 0.0 | 0.0 |
| *Keratella americana* Carlin, 1943 | 201.8 | 451.3 | 9407.8 | 9309.3 | 11.2 | 25.8 | 79.2 | 221.1 |
| *Keratella cochlearis* Gosse, 1851 | 4.9 | 20.2 | 0.2 | 0.8 | 18.8 | 76.8 | 71.2 | 290.9 |
| *Keratella cruciformis* Thompson, 1892 | 0.0 | 0.0 | 29.4 | 121.3 | 0.0 | 0.0 | 0.0 | 0.0 |
| *Keratella lenzi* Hauer, 1953 | 22.3 | 66.0 | 28.6 | 97.8 | 19.6 | 80.8 | 0.0 | 0.0 |
| *Keratella tropica* Apstein, 1907 | 51.5 | 99.4 | 0.0 | 0.0 | 0.0 | 0.0 | 0.0 | 0.0 |
| *Lecane bulla* Gosse, 1851 | 0.2 | 0.8 | 0.0 | 0.0 | 0.0 | 0.0 | 39.2 | 161.7 |
| *Lecane curvicornis* Murray, 1913 | 0.0 | 0.0 | 0.0 | 0.0 | 148.3 | 269.6 | 0.0 | 0.0 |
| *Lecane elsa* Hauer, 1931 | 0.0 | 0.0 | 9.8 | 40.4 | 0.0 | 0.0 | 66.7 | 205.5 |
| *Lecane leontina* Turner, 1892 | 0.0 | 0.0 | 0.0 | 0.0 | 159.3 | 192.3 | 0.0 | 0.0 |
| *Lecane luna* Müller, 1776 | 0.0 | 0.0 | 4.9 | 20.2 | 6.7 | 20.8 | 166.7 | 311.8 |
| *Lecane lunaris Ehrenberg, 1832* | 7.4 | 22.0 | 40.4 | 117.9 | 0.0 | 0.0 | 0.0 | 0.0 |
| *Lecane monostyla Daday, 1897* | 0.0 | 0.0 | 0.0 | 0.0 | 41.1 | 63.8 | 0.0 | 0.0 |
| *Lecane niothis* Harring & Myers, 1926 | 0.0 | 0.0 | 0.0 | 0.0 | 0.2 | 0.8 | 0.0 | 0.0 |
| *Lecane proiecta* Hauer, 1956 | 27.2 | 67.3 | 2674.5 | 6147.7 | 7.8 | 32.3 | 28143.1 | 42661.0 |
| *Lecane scutata* Harring & Myers, 1926 | 39.2 | 161.7 | 0.0 | 0.0 | 0.0 | 0.0 | 0.0 | 0.0 |
| *Lecane signifera* Jennings, 1896 | 0.2 | 0.8 | 0.0 | 0.0 | 0.9 | 3.6 | 0.2 | 0.8 |
| *Lecane ungulata* Gosse, 1887 | 14.7 | 60.6 | 0.0 | 0.0 | 0.0 | 0.0 | 0.0 | 0.0 |
| *Lepadella astacicola* Hauer, 1926 | 14.7 | 60.6 | 0.0 | 0.0 | 0.0 | 0.0 | 0.0 | 0.0 |
| *Lepadella patella Müller, 1773* | 37.3 | 105.3 | 4173.5 | 6930.0 | 72.5 | 104.6 | 20.0 | 80.8 |
| *Lepadella quadricarinata* Stenroos, 1898 | 0.0 | 0.0 | 0.0 | 0.0 | 1.3 | 3.9 | 0.0 | 0.0 |
| *Lepadella* sp. Bory de St. Vincent, 1822 | 0.0 | 0.0 | 0.0 | 0.0 | 0.6 | 2.4 | 0.0 | 0.0 |
| *Liliferotrocha subtilis Rodewald, 1940* | 7.4 | 30.3 | 47.1 | 194.0 | 117.6 | 485.1 | 0.0 | 0.0 |
| *Macrochaetus sericus* Thorpe, 1893 | 0.0 | 0.0 | 0.0 | 0.0 | 0.2 | 0.8 | 24.5 | 101.1 |
| *Notommata* sp. Ehrenberg, 1830 | 0.0 | 0.0 | 0.0 | 0.0 | 0.0 | 0.0 | 24.5 | 101.1 |
| *Paradicranophorus hudsoni Glascott, 1893* | 0.0 | 0.0 | 0.0 | 0.0 | 0.2 | 0.8 | 0.0 | 0.0 |
| *Plationus patulus macracanthus* Daday, 1905 | 127.8 | 226.8 | 29.4 | 121.3 | 32.3 | 84.0 | 0.0 | 0.0 |
| *Platyias quadricornis Ehrenberg, 1832* | 0.0 | 0.0 | 0.2 | 0.8 | 205.4 | 345.3 | 0.0 | 0.0 |
| *Polyarthra dolichoptera Idelson, 1925* | 0.0 | 0.0 | 0.0 | 0.0 | 2.5 | 10.1 | 0.0 | 0.0 |

| | | | | | | | | | |
|---|---|---|---|---|---|---|---|---|---|
| | *Polyarthra vulgaris Carlin, 1943* | 14.9 | 60.6 | 1616.5 | 3984.4 | 131.2 | 261.0 | 0.0 | 0.0 |
| | *Pompholyx* sp. Gosse, 1851 | 0.0 | 0.0 | 0.0 | 0.0 | 0.0 | 0.0 | 313.7 | 1293.5 |
| | *Proales* cf. *commutata Althaus, 1957* | 0.0 | 0.0 | 0.0 | 0.0 | 9.8 | 40.4 | 0.0 | 0.0 |
| | *Proales* sp. Gosse, 1886 | 0.0 | 0.0 | 0.0 | 0.0 | 7.0 | 28.8 | 24.5 | 101.1 |
| | *Proalides tentaculatus Beauchamp, 1907* | 117.6 | 422.7 | 0.0 | 0.0 | 0.0 | 0.0 | 0.0 | 0.0 |
| | *Ptygura spongicola Bērziņš, 1950* | 14.7 | 60.6 | 0.0 | 0.0 | 0.0 | 0.0 | 0.0 | 0.0 |
| | *Squatinella lamellaris Müller, 1786* | 0.0 | 0.0 | 29.4 | 121.3 | 0.0 | 0.0 | 0.0 | 0.0 |
| | *Synchaeta asymmetrica Koch-Althaus, 1963* | 0.0 | 0.0 | 0.0 | 0.0 | 24.5 | 101.1 | 0.0 | 0.0 |
| | *Synchaeta neopolitana Rousselet, 1902* | 0.0 | 0.0 | 0.0 | 0.0 | 122.7 | 484.2 | 0.0 | 0.0 |
| | *Synchaeta oblonga Ehrenberg, 1832* | 0.0 | 0.0 | 0.0 | 0.0 | 619.6 | 2420.7 | 0.0 | 0.0 |
| | *Synchaeta pectinata Ehrenberg, 1832* | 0.0 | 0.0 | 0.0 | 0.0 | 4.4 | 18.2 | 0.0 | 0.0 |
| | *Testudinella patina Hermann, 1783* | 85.8 | 169.6 | 34.5 | 121.6 | 100.8 | 131.0 | 0.0 | 0.0 |
| | *Trichocerca bicristata Gosse, 1887* | 5.1 | 20.2 | 0.0 | 0.0 | 88.7 | 176.3 | 0.0 | 0.0 |
| | *Trichocerca bidens Lucks, 1912* | 0.0 | 0.0 | 14.7 | 60.6 | 18.6 | 76.8 | 436.7 | 944.2 |
| | *Trichocerca cylindrica Imhof, 1891* | 0.0 | 0.0 | 159.8 | 275.3 | 39.4 | 161.6 | 0.0 | 0.0 |
| | *Trichocerca iernis Gosse, 1887* | 402.7 | 793.4 | 2652.9 | 2424.3 | 28.9 | 57.9 | 186.9 | 506.3 |
| | *Trichocerca longiseta Schrank, 1802* | 0.0 | 0.0 | 0.0 | 0.0 | 1.6 | 6.5 | 0.0 | 0.0 |
| | *Trichocerca marina Daday, 1890* | 0.0 | 0.0 | 0.0 | 0.0 | 0.8 | 3.2 | 0.0 | 0.0 |
| | *Trichocerca* sp. Lamarck, 1801 | 0.0 | 0.0 | 0.0 | 0.0 | 0.0 | 0.0 | 39.2 | 161.7 |
| | *Trichotria cornuta Myers, 1938* | 270.6 | 1090.2 | 0.0 | 0.0 | 6.6 | 27.3 | 0.0 | 0.0 |
| | *Trichotria tetractis Ehrenberg, 1830* | 0.0 | 0.0 | 0.0 | 0.0 | 1.7 | 4.7 | 0.0 | 0.0 |
| | *Trochosphaera aequatorialis Semper, 1872* | 0.0 | 0.0 | 0.0 | 0.0 | 4.3 | 16.1 | 0.0 | 0.0 |
| | *Wierzejskiella elongata Glascott, 1893* | 0.0 | 0.0 | 0.0 | 0.0 | 44.1 | 161.7 | 0.0 | 0.0 |
| Testate amoebae | *Arcella conica Playfair, 1918* | 0.0 | 0.0 | 0.0 | 0.0 | 0.2 | 0.8 | 0.0 | 0.0 |
| | *Arcella costata Ehrenberg, 1847* | 0.0 | 0.0 | 0.0 | 0.0 | 14.6 | 40.9 | 0.0 | 0.0 |
| | *Arcella discoides Ehrenberg, 1843* | 0.0 | 0.0 | 0.0 | 0.0 | 0.0 | 0.0 | 0.2 | 0.8 |
| | *Arcella gibbosa Penard, 1890* | 14.7 | 60.6 | 44.1 | 132.1 | 21.8 | 80.8 | 0.0 | 0.0 |
| | *Arcella hemisphaerica Perty, 1852* | 0.0 | 0.0 | 0.0 | 0.0 | 11.7 | 32.9 | 0.0 | 0.0 |
| | *Arcella megastoma Penard, 1902* | 2.9 | 12.1 | 0.0 | 0.0 | 19.2 | 76.7 | 0.0 | 0.0 |
| | *Arcella mitrata Leidy, 1876* | 0.0 | 0.0 | 0.0 | 0.0 | 2.2 | 9.1 | 0.0 | 0.0 |

| | | | | | | | | |
|---|---|---|---|---|---|---|---|---|
| *Arcella rotundata Playfair, 1918* | 0.0 | 0.0 | 0.0 | 0.0 | 2.2 | 9.1 | 0.0 | 0.0 |
| *Arcella vulgaris Ehrenberg, 1830* | 106.0 | 197.4 | 0.2 | 0.8 | 96.7 | 238.5 | 0.2 | 0.8 |
| *Centropyxis aculeata Ehrenberg, 1838* | 0.0 | 0.0 | 0.0 | 0.0 | 36.4 | 90.0 | 82.7 | 193.5 |
| *Centropyxis arcelloides Penard, 1902* | 0.0 | 0.0 | 0.0 | 0.0 | 0.0 | 0.0 | 39.2 | 161.7 |
| *Centropyxis cassis Wallich, 1864* | 0.0 | 0.0 | 0.0 | 0.0 | 121.2 | 443.6 | 0.0 | 0.0 |
| *Centropyxis discoides Penard, 1902* | 4.9 | 20.2 | 0.0 | 0.0 | 19.6 | 80.8 | 23.5 | 97.0 |
| *Centropyxis ecornis Ehrenberg, 1841* | 0.0 | 0.0 | 0.0 | 0.0 | 0.0 | 0.0 | 0.7 | 2.9 |
| *Centropyxis gibba Deflandre, 1929* | 3.1 | 12.1 | 41.0 | 117.7 | 0.0 | 0.0 | 0.0 | 0.0 |
| *Centropyxis aculeata* var. *spinosa Cash, 1905* | 0.0 | 0.0 | 0.0 | 0.0 | 0.0 | 0.0 | 23.5 | 97.0 |
| *Cucurbitella dentata LeClerc, 1815* | 0.0 | 0.0 | 0.0 | 0.0 | 110.2 | 401.9 | 0.0 | 0.0 |
| *Cucurbitella mespiliformis Penard, 1902* | 0.2 | 0.8 | 428.4 | 783.4 | 5.3 | 21.8 | 0.0 | 0.0 |
| *Cucurbitella* sp. Penard, 1902 | 14.7 | 60.6 | 0.0 | 0.0 | 0.0 | 0.0 | 0.0 | 0.0 |
| *Difflugia* cf. *penardi Hopkinson, 1909* | 0.0 | 0.0 | 0.0 | 0.0 | 0.0 | 0.0 | 62.7 | 183.3 |
| *Difflugia cylindrus Thomas, 1953; Ogden, 1983* | 0.0 | 0.0 | 0.0 | 0.0 | 2.2 | 9.1 | 0.0 | 0.0 |
| *Difflugia difficilis Thomas, 1954* | 657.2 | 1162.1 | 58.8 | 242.5 | 9.1 | 28.5 | 19.6 | 80.8 |
| *Difflugia elegans Penard, 1890* | 2.5 | 10.1 | 0.0 | 0.0 | 52.2 | 163.7 | 0.2 | 0.8 |
| *Netzelia gramen Penard, 1902; Gomaa et al., 2017* | 0.0 | 0.0 | 0.0 | 0.0 | 0.0 | 0.0 | 20.2 | 80.7 |
| *Difflugia kempnyi Stepanek, 1953* | 0.0 | 0.0 | 0.0 | 0.0 | 7.2 | 27.2 | 0.0 | 0.0 |
| *Difflugia limnetica Levander, 1900; Penard, 1902* | 0.0 | 0.0 | 0.0 | 0.0 | 4.4 | 18.2 | 0.4 | 1.1 |
| *Difflugia lobostoma Leidy, 1879* | 336.2 | 799.5 | 1305.9 | 3467.1 | 18.4 | 72.6 | 24.9 | 101.0 |
| *Difflugia oblonga Ehrenberg, 1838* | 0.0 | 0.0 | 0.0 | 0.0 | 22.4 | 59.6 | 0.0 | 0.0 |
| *Difflugia pleustonica Dioni, 1970* | 0.0 | 0.0 | 0.0 | 0.0 | 6.1 | 25.3 | 0.0 | 0.0 |
| *Difflugia* sp. Leclerc, 1815 | 31.9 | 121.1 | 0.0 | 0.0 | 0.0 | 0.0 | 0.0 | 0.0 |
| *Difflugia urceolata Carter, 1864* | 0.0 | 0.0 | 0.0 | 0.0 | 29.0 | 55.8 | 0.0 | 0.0 |
| *Euglypha filifera Penard, 1890* | 0.0 | 0.0 | 0.0 | 0.0 | 2.2 | 9.1 | 0.0 | 0.0 |
| *Lesquereusia globulosa Thomas et Gauthier-Lièvre, 1959* | 1819.6 | 2824.7 | 0.0 | 0.0 | 0.0 | 0.0 | 0.0 | 0.0 |
| *Lesquereusia* sp. Schlumberger (1845) | 0.0 | 0.0 | 0.0 | 0.0 | 0.0 | 0.0 | 0.2 | 0.8 |
| *Lesquereusia spiralis Ehrenberg, 1840* | 551.7 | 928.5 | 809.0 | 1211.0 | 174.2 | 382.9 | 0.0 | 0.0 |
| *Nebela collaris Ehrenberg, 1848 sensu Kosakyan et Gomaa, 2013* | 0.0 | 0.0 | 4.9 | 20.2 | 0.0 | 0.0 | 0.0 | 0.0 |

| | | | | | | | | |
|---|---|---|---|---|---|---|---|---|
| *Netzelia corona Wallich, 1864; Gooma et al., 2017* | 0.0 | 0.0 | 0.0 | 0.0 | 0.0 | 0.0 | 0.4 | 1.6 |
| *Netzelia labeosa* Beyens & Chardez, 1997, | 58.8 | 242.5 | 0.0 | 0.0 | 0.0 | 0.0 | 0.0 | 0.0 |
| *Netzelia muriformis Gauthier-Lièvre et Thomas, 1958 new comb.* | 0.0 | 0.0 | 0.0 | 0.0 | 2.2 | 9.1 | 0.0 | 0.0 |
| *Netzelia oviformis Cash, 1909; Ogden, 1979* | 44.1 | 181.9 | 47.1 | 194.0 | 0.0 | 0.0 | 0.0 | 0.0 |
| *Netzelia tuberculata (Wallich, 1864)* | 39.4 | 161.6 | 313.7 | 899.2 | 1.2 | 4.1 | 4.9 | 20.2 |
| *Padaungiella tubulata Brown, 191; Lara et Todorov, 2012* | 0.0 | 0.0 | 25.5 | 105.1 | 0.0 | 0.0 | 0.0 | 0.0 |
| *Sphenoderia lenta Schlumberger, 1845* | 78.4 | 323.4 | 0.0 | 0.0 | 0.0 | 0.0 | 0.0 | 0.0 |
| *Trinema enchelys Ehrenberg, 1838* | 4.9 | 20.2 | 0.0 | 0.0 | 0.0 | 0.0 | 0.0 | 0.0 |
| *Trinema lineare Penard, 1890* | 395.1 | 1325.6 | 4962.7 | 11159.8 | 215.7 | 889.3 | 0.0 | 0.0 |
| *Trinema* sp. Dujardin, 1841 | 0.0 | 0.0 | 29.4 | 121.3 | 0.2 | 0.8 | 0.0 | 0.0 |

**Table S2** – Environmental limnologic variables for hydrological period. SD= Standard Deviation.

| Variables | Flooding | | | Flushing | | | High water | | | Low water | | |
|---|---|---|---|---|---|---|---|---|---|---|---|---|
| | Mean | SD | CV (%) | Mean | SD | CV (%) | Mean | SD | CV (%) | Mean | SD | CV (%) |
| Alkalinity (mg/L) | 19.59 | 4.09 | 20.88 | 12.99 | 1.83 | 14.12 | 18.10 | 2.21 | 12.23 | 13.55 | 6.86 | 50.61 |
| Ammonia (mg/L) | 0.04 | 0.04 | 109.48 | 0.03 | 0.05 | 174.41 | 0.06 | 0.05 | 86.88 | 0.21 | 0.12 | 56.83 |
| Blue-green algae (µg/L) | 0.23 | 0.35 | 154.46 | 1.77 | 1.40 | 79.22 | 0.08 | 0.07 | 88.96 | 3.68 | 2.66 | 72.29 |
| Conductivity (µS/cm) | 70.29 | 12.93 | 18.40 | 46.76 | 4.70 | 10.04 | 43.00 | 3.20 | 7.45 | 49.12 | 14.10 | 28.71 |
| Dissolved Oxygen (mg/L) | 6.25 | 0.93 | 14.96 | 6.59 | 3.37 | 51.21 | 4.02 | 0.97 | 24.21 | 7.59 | 0.79 | 10.47 |
| Fluorescent dissolved organic matter (raw) | 11.16 | 3.06 | 27.38 | 34.64 | 40.78 | 117.72 | 16.48 | 0.82 | 4.97 | 7.08 | 3.27 | 46.14 |
| Nitrate (mg/L) | 0.07 | 0.04 | 62.21 | 0.07 | 0.06 | 88.31 | 0.08 | 0.03 | 42.81 | 0.08 | 0.10 | 121.53 |
| pH | 7.17 | 0.32 | 4.50 | 7.62 | 1.00 | 13.18 | 6.63 | 0.11 | 1.69 | 7.84 | 0.72 | 9.24 |
| Silica (mg/L) | 2.43 | 0.38 | 15.75 | 2.92 | 0.34 | 11.68 | 2.60 | 0.42 | 16.10 | 3.61 | 0.75 | 20.69 |
| Temperature (ºC) | 31.01 | 0.82 | 2.64 | 31.30 | 0.99 | 3.16 | 30.08 | 0.63 | 2.08 | 31.51 | 0.84 | 2.65 |
| Total chlorophyll (µg/L) | 3.59 | 1.07 | 29.74 | 6.57 | 2.27 | 34.59 | 3.99 | 1.43 | 35.96 | 10.86 | 3.12 | 28.70 |
| Total dissolved nitrogen (mg/L) | 0.22 | 0.08 | 35.60 | 0.28 | 0.08 | 30.30 | 0.27 | 0.05 | 18.61 | 0.33 | 0.09 | 25.62 |
| Total dissolved solids (mg/L) | 45.71 | 8.25 | 18.05 | 30.47 | 3.06 | 10.06 | 28.00 | 2.09 | 7.47 | 31.88 | 9.21 | 28.87 |
| Total nitrogen (mg/L) | 0.40 | 0.10 | 24.65 | 0.30 | 0.10 | 34.00 | 0.36 | 0.07 | 19.12 | 0.44 | 0.11 | 25.68 |
| Total phosphorus (mg/L) | 0.09 | 0.04 | 41.79 | 0.05 | 0.03 | 52.75 | 0.06 | 0.02 | 30.33 | 0.05 | 0.02 | 48.47 |
| Turbidity (NTU) | 20.36 | 6.39 | 31.40 | 22.82 | 11.50 | 50.37 | 7.09 | 3.35 | 47.27 | 47.38 | 19.31 | 40.76 |

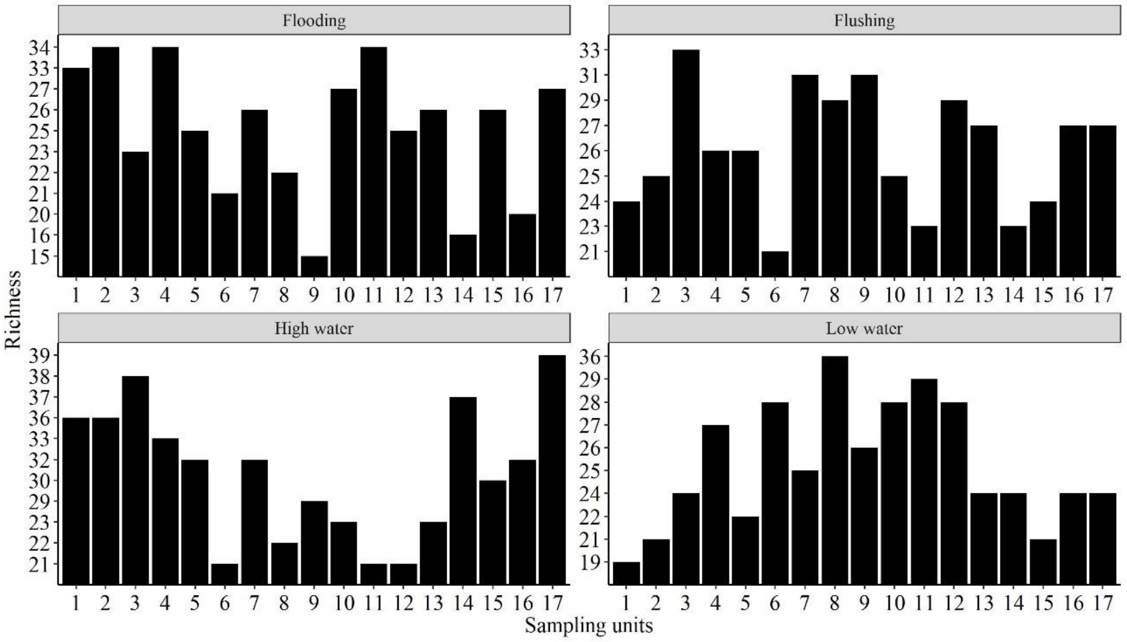


**Figure S1.** Zooplankton species richness per sampling unit in each hydrological period

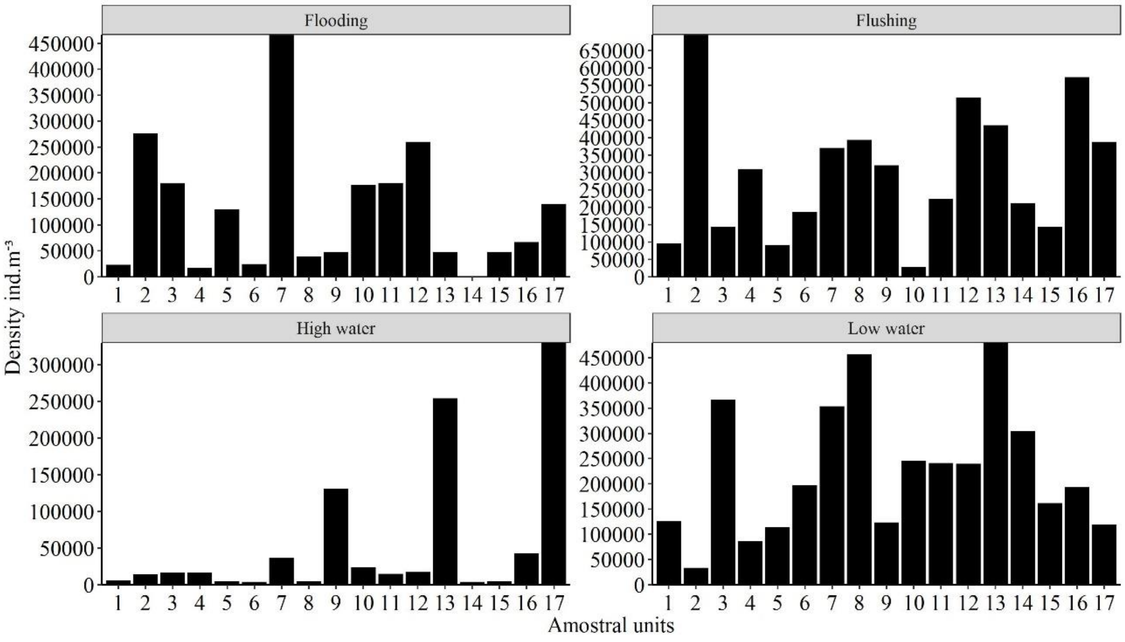


**Figure S2.** Zooplankton density per sampling unit in each hydrological period
